# NetFlow: A Framework to Explore Topological Representations of High-Dimensional Biomedical Data

**DOI:** 10.1101/2025.10.24.683878

**Authors:** Rena Elkin, Jung Hun Oh, Anish K. Simhal, Joseph O. Deasy

## Abstract

Biomedical datasets are increasingly high-dimensional and multi-modal, theoretically enabling a more comprehensive understanding of biological systems. However, common clustering and dimensionality reduction methods often oversimplify continua and obscure complex, biologically relevant patterns, hindering our ability to meaningfully probe sample relationships. To address this challenge, we developed NetFlow, a computational framework that constructs graph representations of structural relationships between samples by integrating a data-driven, lineage-tracing-inspired pseudo-ordering with sparse similarity networks. NetFlow is suitable for small cohorts, features a flexible, modular design, supports multi-modal data fusion, and provides interactive visualization, enabling nuanced exploration of heterogeneity across biological samples. Applied across uni- and multi-modal cancer datasets, NetFlow refined clinically relevant subtypes, identified biomarker associations, and revealed more informative and structured relational patterns when benchmarked against current approaches. NetFlow thus delivers an interpretable graph representation of sample relatedness that supports precision oncology, hypothesis generation, and general high-dimensional data analysis.

## 1 Introduction

High-dimensional bioassays across multiple modalities are collected to study tumor biology, evolution, therapeutic response, and patient prognosis. A central challenge is maximizing the scientific value of this information. This includes modeling and visualizing sample relatedness for meaningful data exploration and hypothesis generation. While clustering and dimensionality reduction techniques, such as PCA [1], diffusion maps [2], t-SNE [3], and UMAP [4], help reveal meaningful patterns in high-dimensional data, they typically perform best when data exhibit strong clustering and may fail to represent continuous variations or partial relationships between subtypes and across data modalities. Biomedical data often fall along a continuum and contain heterogeneity within and across subgroups, which pose challenges for these paradigms. Even when strong clusters are present, visualization often oversimplifies the data into a highly reduced number of dimensions, suppressing nuanced patterns. Topological data analysis (TDA), particularly the Mapper algorithm [5], provides a network-based alternative for capturing global and local structure that preserves topological features of the data, including continuous variation. However, Mapper requires a heuristic selection of a filter function for ordering samples, and importantly, yields overlapping sample clusters that hinder interpretability. Thus, with current approaches, it is still challenging to simultaneously capture clustering and progression along biological continua in an interpretable manner.

Here, we introduce NetFlow, an unsupervised framework for constructing reproducible sample networks that capture both clustering and ordered variation within and between subgroups. NetFlow builds a topological representation we refer to as the Pseudo-Organizational StructurE (POSE), derived from the underlying metric structure of the data using lineage tracing-inspired, data-driven, pseudo-ordering and local similarity networks. Networks offer a natural, human-visible means to investigate how samples organize. They yield interpretable visualizations, as networks are typically less visually cluttered than heatmaps, and their edges provide additional relational information between data points. Moreover, The POSE does not require any assumptions for a low-dimensional embedding, making it well-suited for investigating complex and heterogeneous biological datasets, even with small sample sizes.

We demonstrate the versatility of NetFlow across diverse uni- and multi-modal datasets spanning four cancer types. Use-cases include integrated multi-omics breast cancer (BC) analysis, investigation of *WEE1* as a biomarker in multiple myeloma (MM) using RNA-Seq data, fusion of radiomic feature networks in non-small cell lung cancer (NSCLC), and a fully multi-modal fusion of glioblastoma multiforme (GBM) multi-omics data. We validate NetFlow on simulated data and these case studies, and benchmark it against popular approaches. Our results show that the POSE retains more organizational structure, as reflected by edges that indicate sample similarity and gradual variation in the feature space, compared to visualizations produced by projection and embedding techniques, which inevitably incur a loss of information. We demonstrate that co-localization of samples in the POSE is biologically informative and helps identify subtypes or groupings associated with distinct survival profiles. To facilitate data exploration, we also developed an interactive visualization tool to help researchers investigate their datasets.

## 2 Results

### 2.1 NetFlow Overview

We briefly describe the NetFlow pipeline. Case-specific choices for each use-case are provided individually. Consider an arbitrary dataset *X* = [*x*_1_, *x*_2_, …, *x*_*n*_] ∈ ℝ^*m*×*n*^ with *n* samples and *m* features, where features can be generated and pre-processed in any manner.

NetFlow represents relationships among samples as a graph 𝒢_*POSE*_ = (*V, E*) in an abstract sample-space. Each sample *i* is represented as a vertex *v*_*i*_ ∈ *V* for *i* ∈ {1, …, *n*}. We seek the edges *e* ∈ *E* that capture clustering as well as ordering of the samples based on their feature profiles *x*_*i*_ ∈ ℝ^*m*^. To this end, NetFlow combines a pseudo-ordering backbone that reflects how samples are ordered according to the progression of continuous features, with nearest neighbor connections that reflect clustering based on feature similarities. In this manner, the POSE captures both global and local organizational trends in the data. For a given dataset *X*, the pipeline consists of the following key steps (Fig. 1, Supplementary Algorithm 1):

1. Distance computation: Sample-pairwise distances are computed between feature profiles with respect to a user-selected metric (e.g., Euclidean or optimal mass transport-based Wasserstein metric).
2. Similarity computation: Distances are converted to similarities using kernel functions, resulting in an affinity matrix *K*. This provides a framework for multi-modal and multi-metric analyses.
3. Diffusion-based distances: The affinities are used to model the sample-sample similarities as a weighted graph 𝒢_*MC*_. Multi-scale or scale-free diffusion distances are then computed from the transition matrix *P*. associated with the Markov chain on 𝒢_*MC*_.
4. POSE construction: The global pseudo-ordering backbone is computed via lineage tracing techniques and combined with local similarities captured via edges in the (mutual) kNNgraph to create the POSE network 𝒢_*POSE*_.
5. Visualization: 𝒢_*POSE*_ is visualized, where nodes represent samples and edges indicate relational structure, for user-driven exploratory analysis.

**Fig. 1.**
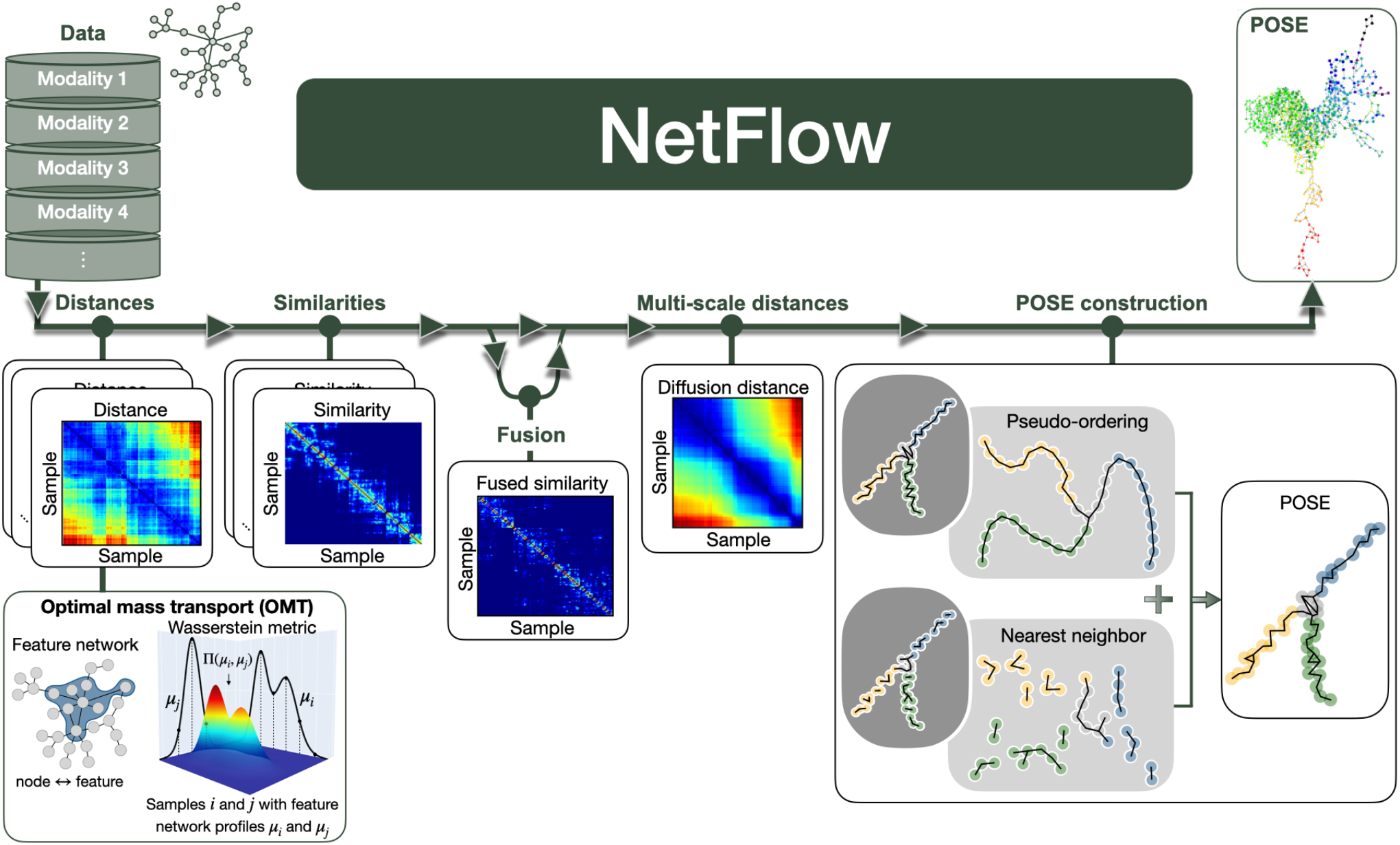
NetFlow analysis pipeline. The input to NetFlow is biomedical feature data and optionally, a feature network. Without any data fitting or machine learning, NetFlow estimates the similarity between all samples and outputs the POSE, a graph representation of the dataset that identifies sample nearness and relatedness. Nodes in the POSE correspond to samples, which are connected by edges based on similarity and ordered variation. Modular components of the NetFlow pipeline, shown from left-to-right, entail computing sample-pairwise distances (e.g., the Wasserstein distance), computing sample-pairwise similarities, optionally fusing similarity structures in the case of multi-modal analysis, computing sample-pairwise multi-scale distances, and computing the pseudo-ordering and nearest neighbor information to construct the final POSE. The POSE is then presented for interactive visualization and user-driven statistical analysis supported within a Jupyter notebook. Visualization may be color-coded by any feature, including genetic, imaging, or survival data, thereby supporting exploratory data analysis.

#### 1. Distance computation

Sample-pairwise distances are computed between feature profiles with respect to a user-selected metric (e.g., Euclidean or optimal mass transport-based Wasserstein metric).

#### 2. Similarity computation

Distances are converted to similarities using kernel functions, resulting in an affinity matrix *K*. This provides a framework for multi-modal and multi-metric analyses.

#### 3. Diffusion-based distances

The affinities are used to model the sample-sample similarities as a weighted graph 𝒢_*MC*_. Multi-scale or scale-free diffusion distances are then computed from the transition matrix *P* associated with the Markov chain on 𝒢_*MC*_.

#### 4. POSE construction

The global pseudo-ordering backbone is computed via lineage tracing techniques and combined with local similarities captured via edges in the (mutual) kNN-graph to create the POSE network 𝒢_*POSE*_.

#### 5. Visualization

𝒢_*POSE*_ is visualized, where nodes represent samples and edges indicate relational structure, for user-driven exploratory analysis.

We provide some insight for each of the key steps in the next few sections. A detailed description is provided in the Methods.

#### 2.1.1 Distance computation

The first step in the NetFlow pipeline is selecting a metric *d* to quantify sample similarity, where *d*_*ij*_ = *d*(*x*_*i*_, *x*_*j*_). The metric will influence which characteristics are reflected in the resulting organizational structure and should therefore be chosen accordingly. The Euclidean metric compares features independently. Whereas the network-based Wasserstein distance [6], a metric for comparing probability distributions by optimizing the transport cost associated with redistributing the initial distribution to match the target distribution, leverages feature interdependencies that may better reflect biological systems such as pathway level interactions or feature correlations. In the context of feature networks, distributions are defined over the set of vertices (i.e., features) and the transport cost of moving a one-unit feature on the graph is taken as the underlying graph distance. Consideration of a feature network and transport distances between samples allows NetFlow to account for complex and coordinated biological features as a system without dimensionality reduction.

#### 2.1.2 Similarity construction

A kernel function *k* is used to construct the similarity structure, *K*_*ij*_ = *k*_*d*_ (*x*_*i*_, *x*_*j*_), from the distances *d*. Kernels inform on geometric structure and help with signal denoising. The similarity structure, called the affinity matrix, implicitly defines a weighted graph where the samples are the vertices and *K*_*ij*_ defines the edge weight between samples *i* and *j*.

#### 2.1.3 Diffusion-based distances

Normalizing the similarity structure by the sample density, *c*_*i*_ = Σ_*j*_*K*_*ij*_, yields the diffusion operator *P* = *D*^−1^*K*, where *D* is a diagonal matrix with entries *D*_*ii*_ = *c*_*i*_. The operator *P* corresponds to a row-stochastic transition matrix that defines a Markov chain on the graph, which models random walks or diffusion processes governed by the sample affinities. This forms the basis of numerous multi-scale and scale-free techniques, such as diffusion maps [2], which have led to important advances in teasing out strong relational signals. We refer to the class of multi-scale or scale-free metrics computed from *P* as diffusion distances. These metrics incorporate the geometry of the underlying space, making them particularly effective at capturing meaningful relationships. While any such metric may be used, we consider the diffusion pseudotime (DPT) metric [7].

#### 2.1.4 Multi-modal integration

Differences in distribution and noise make it challenging to integrate feature data or distances from different modalities, such as imaging and genomics. Various integration techniques have been introduced. Some generate integrated feature profiles for subsequent analysis [8,9], while others analyze each modality independently before integration [10,11]. Either independent or integrated profiles can be used as input feature data for investigation with the NetFlow pipeline. Several methods in the former category treat each modality as a similarity network and perform the integration via the transition matrices [12–14]. We propose integrating the similarities directly. The kernels provide a foundation for comparison by transforming distances from different modalities and metrics onto the same scale ranging from 0 to 1. Averaging these yields what we refer to as a “fused similarity”. Computing diffusion distances on the fused similarity helps control for accumulated modality-specific artifacts and noise. An advantage of multi-scale integration techniques is their ability to control noise, though a key challenge is selecting an appropriate scale. One proposed strategy [15] determines the scale based on the inflection point of the spectral entropy [16] for each modality. Alternatively, scale-free approaches such as DPT [7] circumvent the need to choose a scale altogether. NetFlow’s modular design permits straightforward incorporation of almost any technique for performing the fusion step into its pipeline.

#### 2.1.5 POSE construction

Any distance matrix can serve as input for computing the POSE; however, diffusion-based distances are utilized due to their advantageous geometric and denoising properties. POSE construction begins with inferring the branched organizational backbone topology. This is achieved via lineage tracing, a technique that estimates the pseudo-temporal ordering of cells based on their similarity and likelihood of state transitions, thereby modeling the trajectory a cell takes to reach cell fate [17]. In our framework, we do not view one sample as transitioning into another. Instead, we interpret greater similarity between two samples as indicative of a higher likelihood of shared characteristics, biological markers, and subtypes. Thus, we leverage lineage tracing primarily to establish a pseudo-ordering backbone. Although other approaches may be suitable, we employed the DPT algorithm [7] to construct the backbone for several reasons: it does not impose prior assumptions on the branching structure, it preserves global structure, and it provides a smooth, global pseudo-ordering compared to methods that rely on direct trajectory fitting or minimum spanning trees.

The branched tree-like topology of the backbone reflects lineage tracing assumptions about the nature of state transitions. This ensures a clear ordering and topological connectivity. However, our objective is not to model temporal evolution but rather to capture the relational structure among samples. The tree-like constraint is therefore too restrictive. To address this, we augment the (pseudo-ordering) backbone with local nearest neighbor relationships, thus accommodating loops and deviations from strict hierarchical organization to better reflect underlying data structures. Nearest neighbor information is commonly represented by kNN graphs. A challenge with using kNN graphs is that small values of k can lead to sparse, disconnected graphs, whereas larger values may result in excessively dense graphs, both of which obscure meaningful relationships or trends. We mitigate these issues by supplementing the pseudo-ordering backbone with NN edges (k=1). This approach ensures a connected topology while minimizing excessive density. Restricting to mutual NN edges provides a sparser topology that reduces the risk of spurious associations with outliers (Fig. 6). By combining the pseudo-ordering backbone with (mutual) NN edges, the POSE captures global and local organizational trends in a manner that balances structural simplicity with biological interpretability. See Methods for more details.

The POSE graph provides a visually interpretable representation of how samples organize, where each node corresponds to a sample. NetFlow is implemented in Python and includes an interactive dashboard for exploratory analysis (Supplementary Fig. 1). This interface allows users to perform statistical analyses with customizable visualization options.

### 2.2 NetFlow use-case examples

#### 2.2.1 NetFlow identifies a high-risk group in breast cancer using multi-omics data

The aWCluster algorithm [9] which computes the Wasserstein distance between multi-omics sample profiles was used to cluster TCGA breast cancer (BC) samples (n=726), resulting in the identification of a poor-survival subtype. The multi-omics profiles were computed by integrating the invariant measures associated with Markov processes defined on protein-protein interaction networks weighted by mRNA, copy number alteration (CNA), and methylation data, referred to as integrated measure (IM) profiles. The poor-survival, higher-risk subtype was identified using hierarchical clustering of the Wasserstein distances between the IM profiles. While statistical testing can identify features distinguishing the subtype, the large number of implicated genes complicates meaningful interpretation and visualization via conventional dendrograms and heatmaps.

NetFlow facilitates the investigation of the identified higher-risk subtype by visualizing its organization within a POSE, denoted the BC-POSE. Annotating the BC-POSE with clinical or - omic features clarifies organizational patterns associated with the feature. The branches themselves reveal interpretable biological patterns. For example, branch 1 is associated with high *ERBB2* (a.k.a. *HER2*) IM values (Mann-Whitney U test [MWU] with FDR Benjamini-Hochberg [BH] correction, *P* = 4.24 × 10^−26^). This can be seen by the distribution of IM *ERBB2* values over the BC-POSE shown in Supplementary Fig. 2, highlighting *ERBB2*’s distinct and graded variation over branch 1. Looking at *ERBB2* for each -omic individually shows that *ERBB2* is consistently enriched in branch 1 (MWU test with FDR-BH correction, *P* < 1 × 10^−15^ for all omics) with omic-specific trends that explicate their respective contributions to the IM profile (Supplementary Fig. 2). Unsurprisingly, branch 1 is predominantly HER2-enriched (Supplementary Fig. 3).

**Fig. 2.**
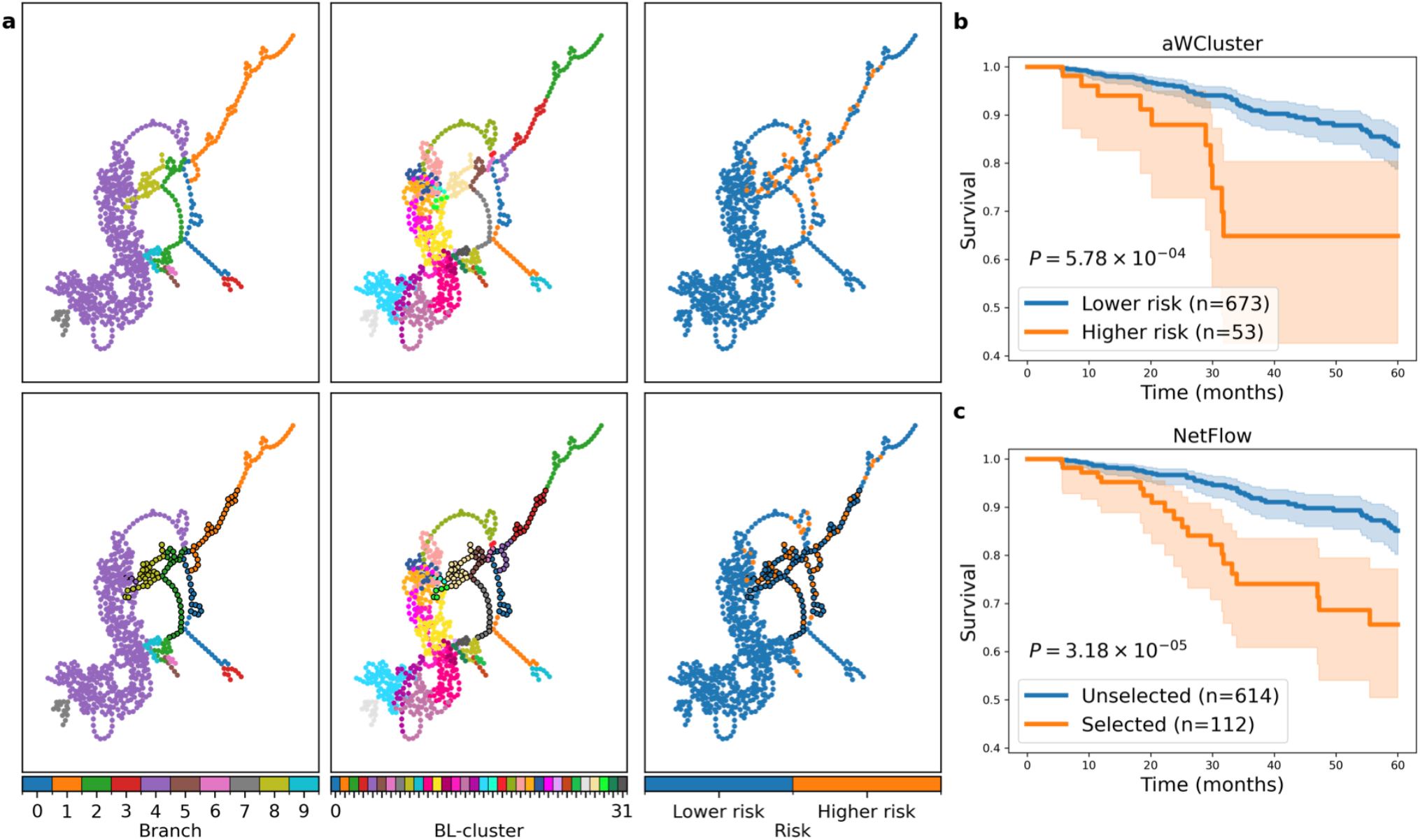
BC-POSE of multi-omics breast cancer (BC) profiles identifies a potential high-risk subtype. (a) The BC-POSE colored respectively from left-to-right by: branch membership, BL-cluster membership, and aWCluster identified risk profile. The same visualization is shown in the bottom row with additional annotation of samples selected from contiguous BL-clusters heavily associated with the aWCluster higher-risk subtype outlined in black. (b) Kaplan-Meier survival curves between higher (orange) and lower (blue) risk subtypes identified by aWCluster (as color-coded in the last column in a). (c) Kaplan-Meier survival curves between NetFlow selected (outlined) and not selected (not outlined) samples (as annotated in the bottom-right subplot in a). (Logrank test *P* values are shown.) Compared to the original clustering method (aWCluster), NetFlow identifies a larger related subgroup of poor survival and related biology.

**Fig. 3.**
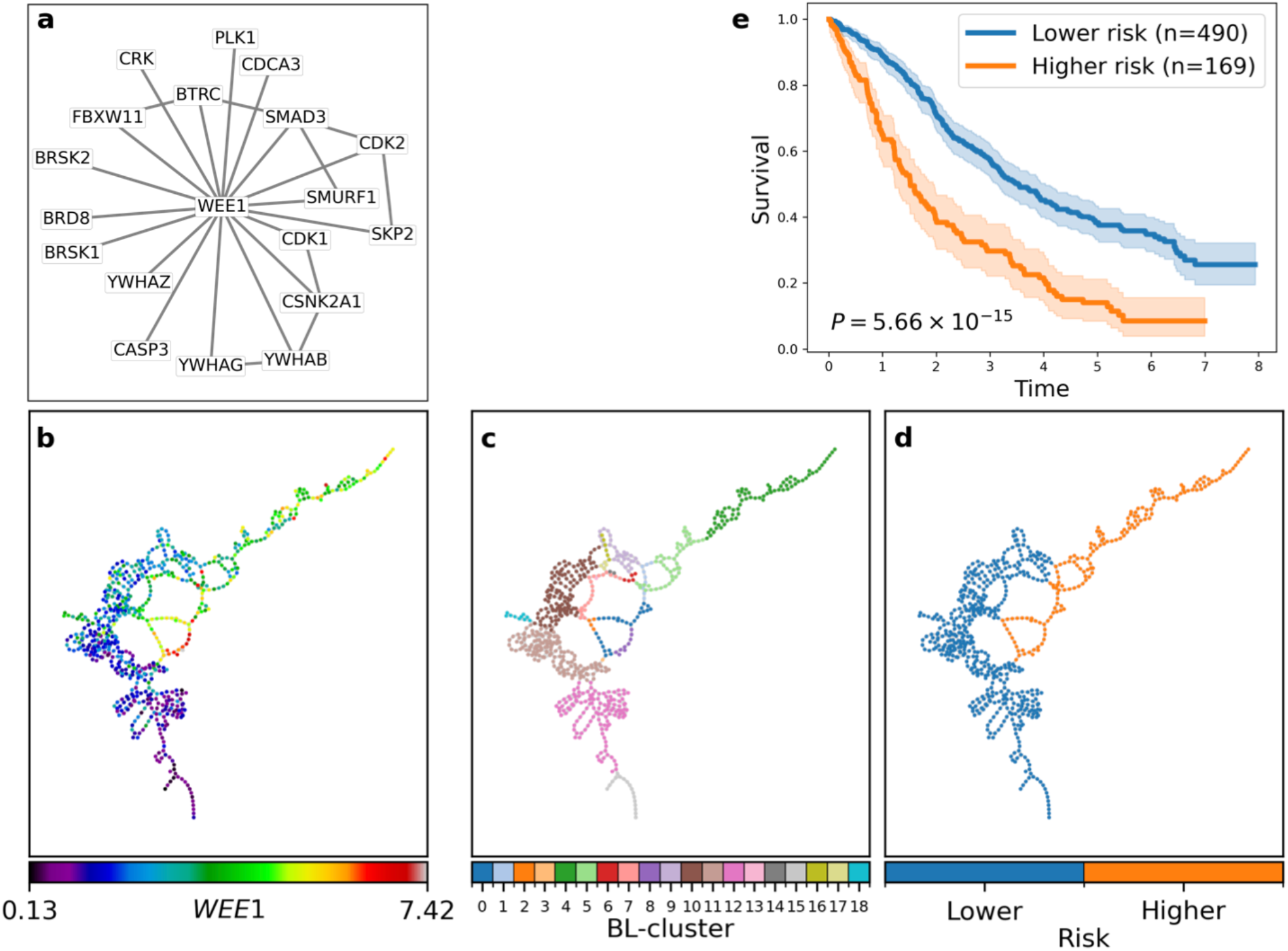
POSE separates multiple myeloma (MM) samples by prognosis based on the *WEE1* gene neighborhood. (a) The *WEE1* neighborhood. (b-c) The MM-POSE based on the scale-free Euclidean distance between *WEE1* neighborhood RNA profiles. Each node represents an individual sample and is colored by: (b) *WEE1* RNA expression, (c) BL-cluster assignment, and (d) NetFlow-defined risk groups. (e) Kaplan-Meier progression-free survival curves (time in years) for the NetFlow identified higher- and lower-risk groups (logrank test *P* value is shown).

Many samples in the aWCluster higher-risk subtype localize near each other in the BC-POSE (Fig. 2a and Supplementary Fig. 4). The exploratory nature of NetFlow supports selecting samples of interest from the POSE. To define subgroups in a more principled manner, we performed Louvain clustering on the POSE and further partitioned the clusters by their intersection with the POSE branches, referred to as BL-clustering (Methods). The branches reflect sequential structure and are therefore not ideal for delineating distinct clusters on their own. However, combining them with the Louvain clusters preserves the ordering, enabling a finer resolution of clustering. Selecting contiguous BL-clusters that predominantly overlap with the aWCluster higher-risk subtype identifies a larger and more statistically robust group (n=112; logrank test, *P* = 3.18 × 10^−5^; Fig. 2c). PAM50 subtype and top multi-omic features that characterize the selected samples are shown in Supplementary Fig. 4 and Supplementary Tables 1-4. Notably, luminal B (LumB) patients within the selected samples show significantly worse survival compared to others (logrank test, *P* = 0.03; Supplementary Fig. 5). Hence, NetFlow facilitates the identification of potential subtypes that share underlying biological characteristics. Importantly, NetFlow does not obscure biological heterogeneity inside or outside the subgroup.

**Fig. 4.**
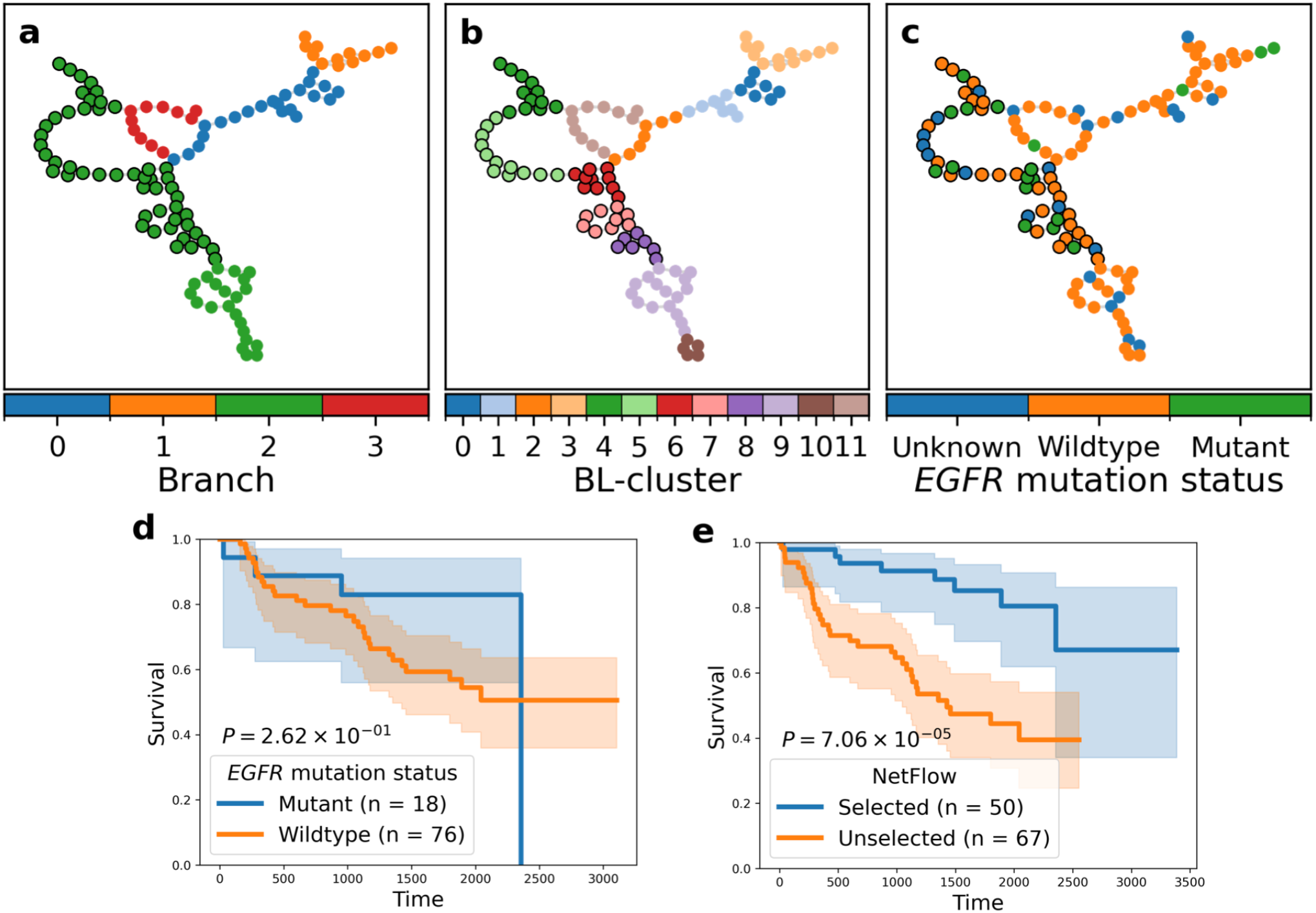
POSE of fused non-small cell lung cancer radiomic network profiles. (a-c) The radio-POSE. Each node represents a sample and the nodes corresponding to selected samples are shown outlined in black. Nodes are colored by (a) branch, (b) BL-cluster, and (c) *EGFR* mutation status. (d-e) Kaplan-Meier survival analysis between (d) *EGFR* mutant vs wildtype samples and (e) NetFlow selected vs not selected samples (logrank test *P* values are shown).

**Fig. 5.**
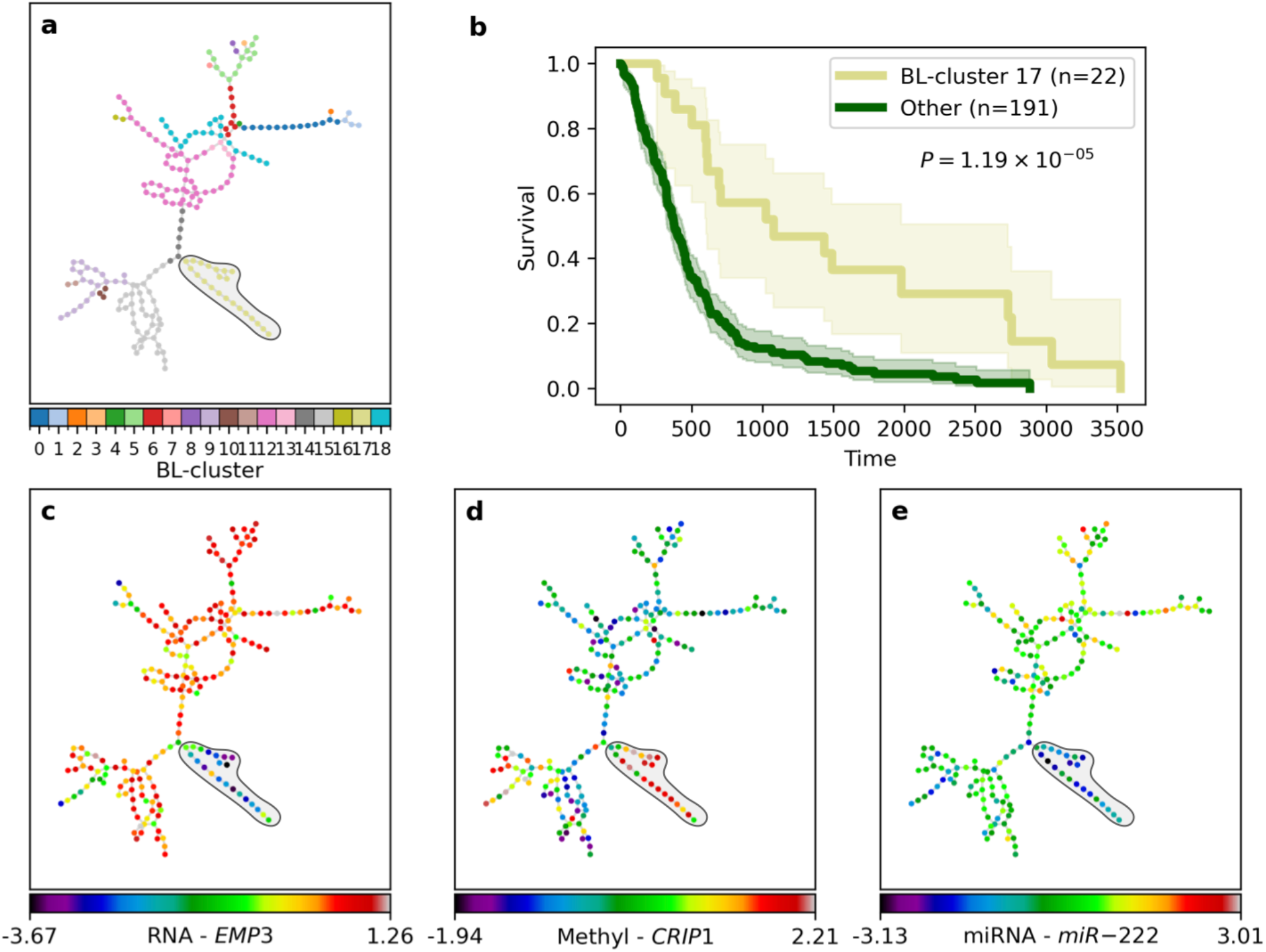
Multi-modal fused POSE reveals a lower-risk subtype in glioblastoma multiforme (GBM). (a) The GBM-POSE. Each sample (i.e., node) is colored by its BL-cluster assignment, accentuating BL-cluster 17. (b) Kaplan-Meier survival analysis between BL-cluster 17 and the rest of the cohort (logrank test *P* value is shown). In (c-e), the GBM-POSE is color-coded by the top features most significantly associated with BL-cluster 17 for each -omic data type, highlighting: (c) low mRNA gene expression of *EMP3*: *P* = 5.77 × 10^−9^, (d) DNA hypermethylation (methyl) of *CRIP1*: *P* = 3.27 × 10^−6^, and (e) low miRNA expression of *miR-222*: *P* = 6.59 × 10^−8^ in BL-cluster 17 (MWU test with FDR-BH correction).

**Fig. 6.**
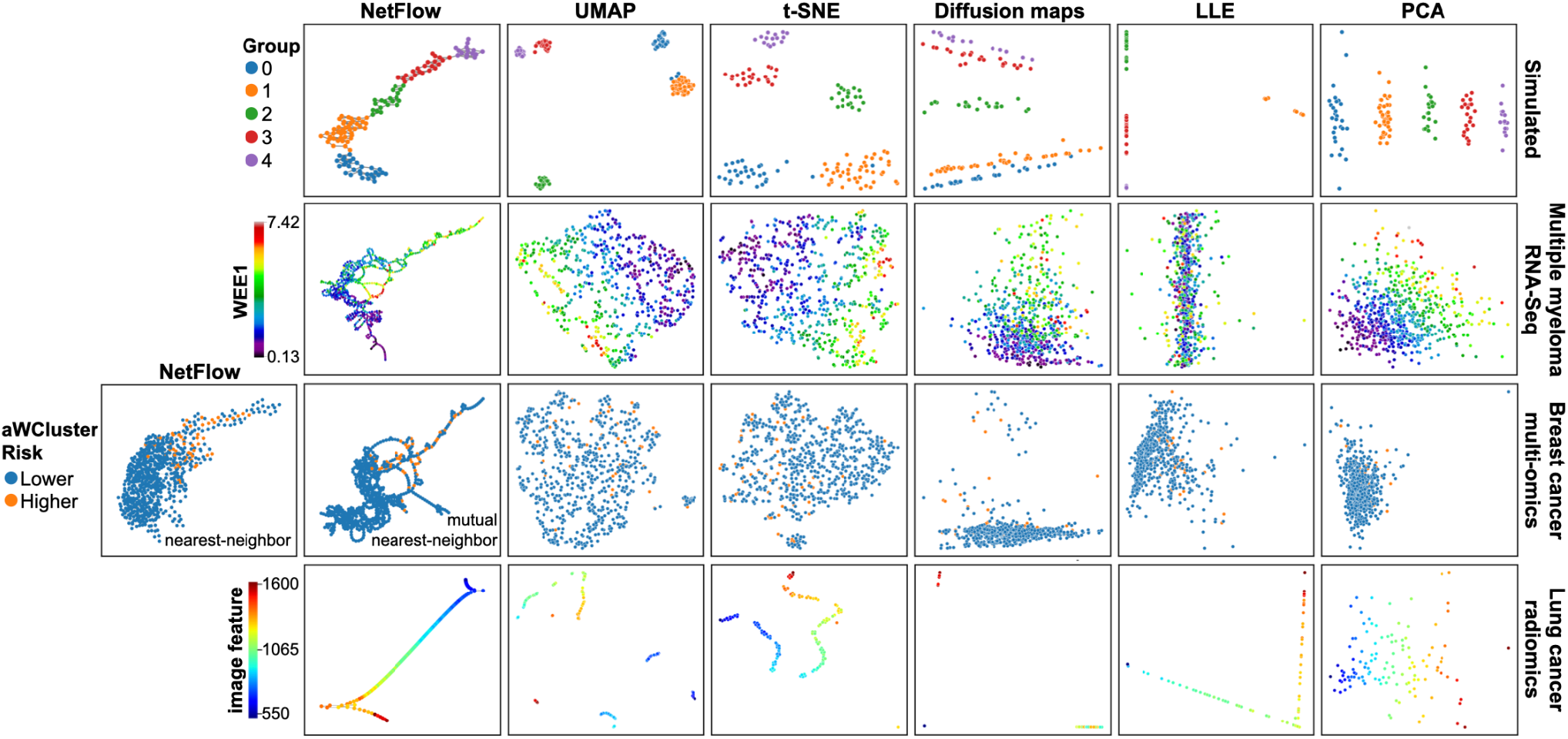
NetFlow performance validation and benchmarking. A comparison of the NetFlow POSE to projected 2-dimensional visualizations of benchmark algorithms for simulated and real-world datasets. Columns correspond to methods and rows reflect datasets. (top row) Simulated data of a cohort comprising five groups generated via the Poisson distribution with increasing mean values according to group. Samples are colored by group membership. (second row) Multiple myeloma RNA-Seq data; samples are colored by *WEE1* expression. (third row) Multi-omics breast cancer integrated aWCluster IM profiles. NetFlow POSEs are shown using (left) nearest neighbors and (right) the default mutual nearest neighbors. Samples are colored by aWCluster identified risk group. (bottom row) Lung cancer radiomics dataset; samples are colored by a Gabor filter feature indicative of edge detection.

#### 2.2.2 NetFlow identifies high-risk cohort in a previously defined WEE1 expression cohort in multiple myeloma

MM remains a cancer without a cure, underscoring the need for further investigation into its genomic underpinnings and trajectories of progression [18]. In a recent study, we identified *WEE1* expression as a prognostic biomarker for identifying higher-risk MM in newly diagnosed patients [19]. Overexpression of *WEE1* presumably increases the time available for DNA repair. To further understand the relationship between the *WEE1* and the set of genes that directly interact with it, we applied the NetFlow pipeline to the Multiple Myeloma Research Foundation’s CoMMpass dataset (IA19) of newly diagnosed MM (NDMM) patients (n=659) [20].

A POSE (MM-POSE) was constructed using the scale-free DPT distance, derived from the Euclidean distance between RNA expression profiles within the *WEE1* gene neighborhood (Fig. 3a), as defined by the Human Protein Reference Database (HPRD) [21]. *WEE1* expression strongly aligns with the inferred sample organization in the MM-POSE (Fig. 3b), affirming that the *WEE1* neighborhood captures meaningful prognostic structure.

We identified distinct higher- and lower-risk groups (Fig. 3d and Supplementary Table 5) by selecting contiguous BL-clusters along the MM-POSE (Fig. 3c). Unlike the prior analysis which examined top and bottom tertiles of patients sorted by *WEE1* expression, our method retains the full cohort, enabling a more complete and statistically significant risk stratification (logrank test, *P* < 1 × 10^−14^; Fig. 3e). The MM-POSE and BL-clustering provides finer-grain subtypes, offering deeper insight into *WEE1*’s biological context (Supplemental Fig. 6). The strong relationship with survival adds to evidence that *WEE1* is an emergent driver of progression in this disease.

#### 2.2.3 Wasserstein-based fused NetFlow analysis identifies association between radiomic and genomic data in lung cancer

Associations between radiomic and genomic features are difficult to resolve, as radiomic features often lack strong correlations with genomic features. NetFlow provides an interactive framework to probe how NSCLC samples organize based on three radiomic feature networks, and whether this structure (radio-POSE) reflects underlying gene expression patterns.

NSCLC computed tomography (CT) imaging data (n=117) were downloaded from The Cancer Imaging Atlas (TCIA) and the matched RNA-Seq gene expression profiles from Gene Expression Omnibus (GEO) database (GSE103584) [22]. Tumor regions of interest (ROIs) were manually contoured by a physician. In total, 397 radiomic features were extracted from ROIs using the CERR software [23] following image biomarker standardization initiative (IBSI) definitions [24]. Radiomic features were broken into strongly correlated networks using partial correlation coefficient analysis. The graphical least absolute shrinkage and selection operator (LASSO) was then applied to eliminate statistically redundant connections, resulting in three connected sub-networks consisting of 68 nodes and 88 edges, 56 nodes and 86 edges, and 17 nodes and 26 edges. The radio-POSE was constructed by fusing the similarity structure associated with each radiomic feature network (Methods, Supplementary Algorithm 2), based on underlying sample-pairwise Wasserstein distances.

The radio-POSE (Fig. 4) was composed agnostic of clinical, mutational, and genomic features, yet we find preferential organizational patterns (Supplementary Fig. 7). Most *EGFR*-mutant samples localized along branch 2 (Fig. 4c). *EGFR* mutation status alone does not significantly stratify survival in this dataset (logrank test, *P* = 0.26; Fig. 4d), but selecting *EGFR*-mutant enriched connected BL-clusters results in significantly stratified survival profiles (logrank test, *P* = 7.06 × 10^−5^; Fig. 4e). We compared RNA expression between selected and unselected samples to inspect the potential relationship between the organizational structure and gene expression. Eight genes were found to be significantly higher for the selected samples (Supplementary Fig. 8; MWU test with FDR-BH correction: *FOLR1* (*P* = 0.01); *CACNB1* (*P* = 0.03); *CTSH* (*P* = 0.04); *CDKL2* (*P* = 0.04); *HNF1B* (*P* = 0.04); *GPC4* (*P* = 0.04); *RAP1GAP* (*P* = 0.04); *C1orf116* (*P* = 0.04)). Notably, *FOLR1* is known to be over-expressed in several solid tumors, particularly lung cancer [25], and has garnered interest as a potential drug target in NSCLC [26]. No such associations were found using the Euclidean distance (Supplementary Fig. 9), highlighting the potential advantage of the Wasserstein metric for comparing samples based on how their features operate within a feature network that may inform on multi-modal associations. Further work is needed to understand these associations, but this application demonstrates the ability of NetFlow to define potential biomarker gene panels, not only for prognosis but also for identifying closely related subtypes.

#### 2.2.4 Multi-omics fused NetFlow analysis identifies a lower-risk subtype in glioblastoma multiforme

To demonstrate NetFlow’s flexibility in integrating multi-modal data, we analyzed TCGA GBM data (n=213) comprising mRNA gene expression, DNA methylation, and microRNA (miRNA) expression profiles. We constructed a multi-omics fused POSE (GBM-POSE; Fig. 5) on pre-processed data from a prior Similarity Network Fusion (SNF) analysis [13] using NetFlow’s multi-modal pipeline (Methods and Supplementary algorithm 2). While SNF was used here for comparison to an established integrative approach, NetFlow’s modular design permits incorporation of SNF or other integration techniques [8,12,14] for performing the fusion step. Subsequent POSE construction offers a powerful representation and visualization for investigating biological implications of sample associations from multi-modal data for any choice of integration technique.

BL-clustering of the GBM-POSE (Fig. 5a) identified a lower-risk subtype (BL-cluster 17) with significantly better survival (logrank test, *P* = 1.19 × 10^−5^; Fig. 5b). This is consistent with the previously reported SNF-defined lower-risk group [13] (Supplementary Fig. 10). A quantitative approach for assessing the relative contribution of each -omic on the edges in the GMB-POSE is described in Supplementary Figs. 11 and 12. Top features associated with BL-cluster 17 (Fig. 5-e and Supplementary Table 6) recapitulate known GBM biology. BL-cluster 17 exhibits significantly lower mRNA expression of *EMP3* and *TIMP1* (MWU test with FDR-BH correction, *P* < 1 × 10^−8^), along with reduced miRNA expression of *miR-222* (MWU test with FDR-BH correction, *P* = 6.59 × 10^−8^). Elevated levels of these markers have been linked to glioma progression, immune modulation, cellular proliferation and resistance to apoptosis [27–30]. These markers are all known to be related to inflammation in GBM, and their downregulation in BL-cluster 17 aligns with its favorable prognosis. While each of these markers has been individually implicated in glioma-related processes, their combined prognostic relevance in GBM remains unexplored. Similarly, the role of *CRIP1*, found to be hypermethylated in BL-cluster 17 (MWU test with FDR-BH correction, *P* = 3.27 × 10^−6^), and its potential regulation via methylation, has not been characterized in the context of GBM.

### 2.3 NetFlow performance and benchmarking

To assess NetFlow’s usefulness, we compare POSE results on simulated and real-world datasets to current approaches, including UMAP, t-SNE, diffusion maps, locally linear embedding (LLE), and principal component analysis (PCA), here referred to as the benchmark methods (Fig. 6).

#### 2.3.1 Simulated dataset

To evaluate NetFlow’s performance, we generated a synthetic dataset using the Poisson distribution (Methods). Subgroups within the cohort (of sizes n = 30, 43, 20, 22, 15) were simulated using increasing mean values to ensure features (m = 500) were drawn from distinct, progressively increasing but overlapping distributions to mimic real-world data, such as bulk RNA-Seq, which typically exhibits subtype-specific variability. Resulting visualizations of the POSE and all benchmark methods depict clustering patterns (Fig. 6; top row). However, UMAP and t-SNE incorrectly group a few samples and critical information regarding relationships between groups is lost. In contrast, the POSE edges encode both local similarity and global organizational structure, capturing meaningful variation across the entire data landscape. As expected, the POSE is seen to capture both clustering and progression of samples and subgroups.

We also considered how variables regarding common characteristics for a given dataset affect the runtime. We used the Poisson distribution to simulate feature data while varying the number of samples, the number of groups, and the number of features (Methods) and compared NetFlow runtime to the benchmark methods (Supplementary Fig. 13). Overall, we find that the number of groups and the number of features have little effect on the runtime across all methods, where t-SNE has the longest runtimes across all iterations and NetFlow runtime tends to be comparable or slightly faster than UMAP. The number of samples has the largest effect on runtime for all approaches, especially NetFlow. NetFlow exhibited runtimes (comparable to Diffusion maps, LLE, and PCA) that were lower than UMAP and t-SNE for low sample sizes. However, once the number of samples exceeds 500, NetFlow runtimes match and then exceed UMAP runtimes. Interestingly, this aligns with the UMAP recommendation of having at least 500 samples, whereupon the benefit in computational efficiency offsets the approximations used in the algorithm [4]. At 1000 samples, NetFlow runtime matches that of t-SNE, after which NetFlow runtime continues to grow at a faster rate than the other methods.

#### 2.3.2 Real-world datasets

We compared NetFlow performance to benchmark methods using the datasets presented in the preceding use-cases (Fig. 6). The BC-POSEs using kNN and mutual kNN edges (k=1) are rendered, demonstrating consistency between them. Using mutual kNN yields a sparser POSE, yet the organization based on aWCluster metrics identifying a higher-risk subtype remains comparable. The MM-POSE based on the *WEE1* neighborhood colored by *WEE1* expression demonstrates the global gradual variation reflected in the POSE (Fig. 3b). Such heterogeneity contributes to the amorphous layout exhibited by benchmark methods, while the POSE retains intra- and inter-cluster organization and reveals transitions. Lastly, since the benchmark methods do not contain an integration step for multi-modal or multi-metric analysis, a single radiomic dataset of 56 features was used for benchmarking the lung cancer data. A Gabor filter edge indicator feature, used to color-code the samples, serves as a representative example showing that the POSE preserves continuous spectra among samples, avoiding the clustering-driven separation of cases typical of other methods. In contrast to the benchmark methods which produce a more unstructured distribution of points, NetFlow consistently yields a finer resolution and graded representation of the relational organization.

## 3 Discussion

NetFlow provides a unified framework for constructing interpretable and reproducible visualizations of sample relationships from high-dimensional data. By combining a pseudo-ordering backbone with local similarity networks, NetFlow preserves both global variation and local clustering in a single, connected topological representation. This approach enables researchers to uncover subtype structures, assess modality contributions, and generate hypotheses from complex data, even with limited sample sizes.

We demonstrated NetFlow applications through diverse case studies. In BC, co-localization in the POSE representation refined a higher-risk subtype previously identified by hierarchical clustering of integrated multi-omics profiles, yielding a biologically meaningful and more statistically significant group. In MM, the POSE resolved a *WEE1*-associated risk group, linking organization with disease free survival in the complete cohort. In NSCLC, the POSE demonstrated survival-based organizational patterns associated with *EGFR* mutation and revealed elusive genomic associations with radiomic-informed structure not captured by Euclidean metrics. Finally, in GBM, the POSE recapitulated an SNF-derived lower-risk subtype using the fused multi-modal pipeline and provided finer resolution for visual clarity. In all these examples, NetFlow’s POSE offered a navigable representation that retained complex relational structure amid the continuity of sample organization, resulting in biological insights.

NetFlow remains well-suited for datasets that exhibit strong, distinct subgroups. Their linkage within the POSE does not imply artificial relationships. Rather, they preserve global inter-group relationships. The POSE edges weighted by the (diffusion) distances retain the structure and strength of these relationships. Coloring nodes by distance to a reference sample or inspecting edge weights enables interpretable visualization of both intra- and inter-group similarity and allows users to assess the degree of relatedness across the network. By leveraging edge weights as a measure of similarity, NetFlow can support decisions such as treatment planning, where outcomes may be expected to align with those of neighboring samples. More broadly, it provides a principled way to assess the appropriateness of sample comparisons by capturing not only which samples are most similar, but also the degree of similarity, which is critical in heterogeneous contexts.

Further development is needed to improve runtime and scalability to larger sample sizes. Future releases will better leverage parallel processing to improve tractability and scalable approximations such as the Sinkhorn algorithm [31] which could be incorporated as an alternative to the powerful but computationally intensive Wasserstein distance. Limitations in the algorithm selected for computing pseudo-ordering should be considered as well. The current work used the DPT algorithm which relies on a connected similarity graph which may underrepresent highly distinct subgroups. We plan to address this in future work with modifications to treat unconnected graphs in the eigen-decomposition [32]. Additionally, although diffusion distances help mitigate effects of noise and outliers, preprocessing to detect and filter outliers remains beneficial.

A key strength of the NetFlow framework is flexibility. NetFlow’s modular design allows for flexible integration with other preprocessing, fusion, or computational methods. NetFlow accommodates diverse data sources through scaled similarity metrics. Benchmarking against standard dimensionality reduction methods further underscored NetFlow’s strengths. While benchmark approaches such as PCA and UMAP provided rough clustering patterns, they often distorted inter-sample distances and failed to capture gradations of variability within and between groups. In contrast, NetFlow preserved both local and global relationships, maintained interpretability through explicit connectivity, and its pipeline supports integration of multi-modality datasets. This makes NetFlow well-suited for data-driven hypothesis generation and exploratory analysis in heterogeneous and complex biological systems.

In conclusion, NetFlow provides a broadly applicable framework for organizing, visualizing, and interpreting high-dimensional biological data. Its POSE representation offers a natural interface for interactive data exploration, stratification, and downstream analysis. By focusing on human-interpretable, graph-based visualizations of similarity, NetFlow brings us closer to a general framework for understanding heterogeneity in biological systems and identifying clinically actionable structure in complex data. Future development will further link relational organization with functional or clinical outcomes and explore supervised extensions to extract biomarkers associated with POSE structure. As datasets continue to grow in size and complexity, tools like NetFlow that prioritize interpretability, flexibility, and biological insight will be essential for translational and integrative data analysis.

## 4 Methods

### 4.1 NetFlow pipeline

Input to NetFlow is uni-or multi-modality data consisting of various features for a set of samples. The samples, e.g., patients, biopsies, tumors, cells, etc., may be any set of observations. The features may be interpretable, e.g., gene expression, methylation, etc., collected from various modalities, but they need not be. For example, the features may have been derived from a data transformation or learned from a deep-learning algorithm applied to the original data. We consider the general dataset with *n* samples and *m* features, denoted *X* ∈ ℝ^*m*×*n*^. For any such input, the NetFlow pipeline can be broadly broken down into the following steps:

1. Compute sample-pairwise distances from the feature dataset.
2. Compute sample-pairwise similarities from distances.
3. (Optional) Fuse similarities across modalities/metrics.
4. Compute diffusion distances (e.g., DPT) from similarities.
5. Compute the POSE from sample-pairwise distances (derived from any of steps 1-4).
6. Visualize and probe the resulting POSE.

The NetFlow framework was intentionally designed in a modular fashion to enhance user interaction with various other pipelines. Each step in the pipeline may be computed within the NetFlow framework and downloaded for use in other analyses. Alternatively, pre-computed data may be uploaded at any step. We therefore describe each step of the NetFlow independently.

#### 4.1.1 Sample-pairwise distances

The NetFlow pipeline starts by computing pairwise-sample distances. A standard metric is the Euclidean distance; however, any metric may be used. The choice of metric determines the aspect of similarity that will be captured to reflect the distances. For example, the Euclidean distance measures point-wise similarity, whereas the Wasserstein distance measures shape similarity.

In general, the 1-Wasserstein distance [33] between probability distributions *μ*_0_ and *μ*_1_ on ℝ^*d*^ is defined as

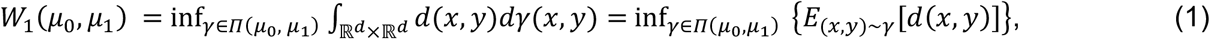

where *Π*(*μ*_0_, *μ*_1_) denotes the set of all couplings on *R*^*d*^ with marginals *μ*_0_ and *μ*_1_, *E*_(*x*,*y*)∼γ_ denotes the expectation given the coupling of random variables *x* ∼ *μ*_0_ and *y* ∼ *μ*_1_ is *γ* and the transportation cost is the distance *d*(*x, y*) = ||*x* − *y*||.

For a simple, connected, undirected graph *G* = (*V, E*) with *m* nodes (|*V*| = *m*) and *r* edges (|*E*| = *r*), the Wasserstein distance between discrete probability distributions *ρ*^0^ and *ρ*^1^ ∈ ℝ^*m*^ defined on the graph *G*, is computed as

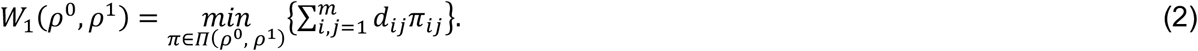

Here, *Π*(*ρ*^0^, *ρ*^1^) denotes the set of all couplings on *G* with marginals *ρ*^0^ and *ρ*^1^, i.e.,

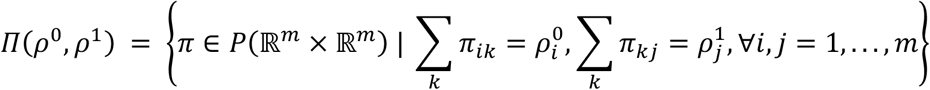

and the transportation cost of moving a unit of mass from node *i* to node *j*, denoted *d*_*ij*_, is the underlying graph distance.

If in addition to a dataset *X* ∈ ℝ^*m*×*n*^, one has a feature network *G*_*X*_(*V, E*) where the nodes *v* ∈ *V* represent each of the *m* features and the edges *e* ∈ *E* indicate relationships between the features (i.e., nodes), sample-pairwise distances may be computed on feature neighborhoods or graph profiles (over the whole graph or on a specified feature subset of interest) via the Wasserstein distance (Eq. 2).

The sample-pairwise distance matrix computed on the feature dataset *X* with metric *M* is denoted 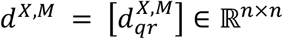, or simply as *d* when clarification is not needed. The distances may be computed with any metric supported in the NetFlow package or uploaded from any pre-computed analysis.

#### 4.1.2 Sample-pairwise similarities

For feature dataset *X* and any choice of metric *M*, the sample-pairwise similarity matrix 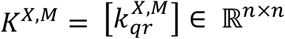, for all samples *q, r*, is constructed using some kernel function *k*. In this work, we use the (adaptive) Gaussian kernel [7]:

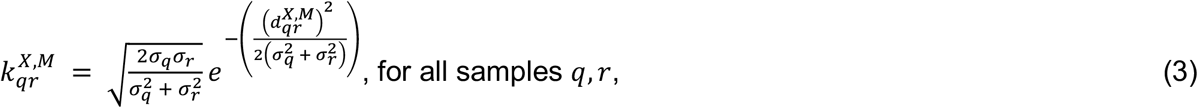

where 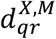 denotes the corresponding distances as previously described. The kernel width *σ*_*q*_ specifies the range of each sample *q*’s reachable neighbors. The user can choose to set *σ*_*q*_ as a constant value for all samples *q*, or use the adaptive kernel and set *σ*_*q*_ as the distance to its k^th^ nearest neighbor for a user-specified choice of k. The latter produces a similarity close to one for distances within the standard deviation of the distance distribution. This is comparable to the kNN graph construction that is the foundation of many dimensional reduction techniques including UMAP, locally linear embedding [34], Isomap [35], laplacian eigenmaps [36], and pseudotime algorithms [7,32]. In contrast to these approaches, here, the similarity structure is not restricted to nearest neighbors. By taking 1 minus *K*, a similarity matrix can be treated as a distance *d*^*K*^:

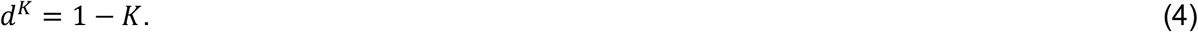

#### 4.1.3 Fused similarities

Differences in the scale and distribution of distances computed via different metrics or on features from different modalities make it challenging to meaningfully combine differential structures. The (adaptive) Gaussian kernel transforms the distances to similarities on the same scale with the range of values constrained to [0, 1]. The resulting similarities have a Gaussian distribution with respect to the chosen metric on the feature space and the adaptive form of the Gaussian kernel provides further standardization by imposing sample-specific normalization. Further considerations regarding the distribution of similarities associated with different feature modalities and metrics are needed. However, the similarities lie in the same scale-space which therefore provides a more natural setting as a starting point to superpose the relational associations and construct a fused similarity matrix. Currently, NetFlow supports computing a fused similarity matrix, denoted 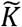, from a set of *s* similarity matrices 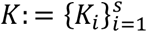 (where the *K*_*i*_’s are computed via various metrics and modalities for the same group of samples), as the linear combination:

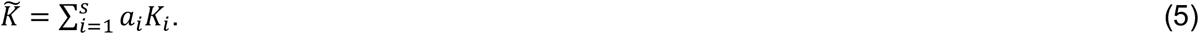

The default behavior sets the weights *a*_*i*_ (*i* = 1, …, *s*) to be uniform (*a*_*i*_ = 1/*s*) unless otherwise specified by the user.

#### 4.1.4 Diffusion distance

Unless specified otherwise, scale-free diffusion distances are computed from the (fused) similarity via the diffusion pseudotime (DPT) metric, as described in [7]. In the context of lineage tracing, diffusion maps [2] provide a basis for the manifold underlying the high-dimensional feature set determined by the multi-scale transition matrix describing the probability of transitioning from one state to another along the differentiation process. The DPT then approximates the geodesic distance between cells on this manifold.

#### 4.1.5 Computing the POSE

For any distance matrix, *d* ∈ ℝ^*n*×*n*^, the POSE is generated in two main steps: First, the branched pseudo-ordering is computed according to the DPT algorithm as described in [7] with adapted code and slight modifications to the SCANPY implementation [37]. Branches are determined by the anti-correlational structure of the diffusion distances between endpoints of a trajectory. Here, the root is assigned as the sample with the largest average distance to all other samples:

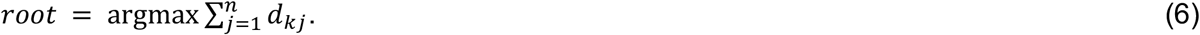

The POSE backbone topology is then constructed by creating a node for each sample. Next, edges are assigned between samples (i.e., nodes) that appear consecutively in each branch of the resulting pseudo-ordering, and between samples that connect the branches. Second, edges between mutual kNNs are added to complete the POSE. Some datasets may justify using NN edges over mutual NNs, so it is offered as an option in the NetFlow library. NN edges for k>1 may be incorporated but the default behavior is to use mutual (k=1) NN edges only.

#### 4.1.6 Probing the POSE

Probing the POSE broadly refers to visual inspection and downstream analyses, such as differential analysis of features, survival, response, etc. between branches, selected co-localized samples, or clusters. The branches reflect global ordering and transitions and therefore do not explicitly represent distinct clusters. Edge distances can be used to assess how strongly neighboring samples are associated with each other, and the proposed BL-clustering can be used to identify clusters.

#### 4.1.7 BL-clustering

Let 𝒮 = {*s*_1_, *s*_2_, …, *s*_*n*_} denote the set of samples and let 𝒢_*POSE*_ = (*V, E*) be the POSE graph, where each node *v*_*i*_ corresponds to a sample *s*_*i*_ ∈ 𝒮. We define the function *β*: 𝒮 → {1,2, …, *B*} that maps the samples to the pseudo-ordering partition of the POSE derived from the lineage tracing procedure into *B* branches so that *β*(*s*_*i*_) = *b* indicates that sample *s*_*i*_ belongs to branch *b*. Similarly, performing Louvain clustering on the POSE, where distances can be used to define edge weights, partitions the samples into *C* clusters assigned via *λ*: 𝒮 → {1,2, …, *C*}. The intersection of branches and clusters defines a refined partition 𝒞_*b*,*c*_ = {*s* ∈ 𝒮 | *β*(*s*) = *b and λ*(*s*) = *c*}, referred to as the BL-clustering. Clustering patterns may be enhanced by considering kNN edges with higher values of k (or alternatively, using a distance weighted cutoff) in the POSE construction.

### 4.2 Data

#### 4.2.1 Breast cancer

RNA, CNA, and methylation profiles for 726 breast cancer samples were taken from TCGA, as described in [9]. The integrated multi-omics profiles and Wasserstein distances were computed as described in [9].

The BC-POSE was constructed as follows: Similarities were computed from the Wasserstein distances using ‘k_nn=12’ as the kernel width (Eq. 3). The DPT distance was then used to construct the POSE using ‘n_branches=3`.

#### 4.2.2 Multiple myeloma

The RNA-seq data was taken from the Multiple Myeloma Research Foundation’s CoMMpass dataset, release version 19. Details on the dataset, collection, and curation methods have been previously published [20]. The inclusion criteria for this study and the preprocessing and feature computations are described in detail in [19,38]. Briefly, 659 subjects with RNA-seq data were analyzed. High-risk *WEE1* was defined as subjects with the top third of *WEE1* expression in the CoMMpass cohort and low-risk *WEE1* was defined as subjects with the bottom third of *WEE1* expression. Gene interaction information was taken from the HPRD [21].

The MM-POSE was constructed as follows: Euclidean distances were computed between *WEE1*-neighborhood expression profiles. Similarities were then computed using ‘k_nn=12’ as the kernel width (Eq. 3). The DPT distance was then used to construct the POSE using ‘n_branches=3`.

#### 4.2.3 Lung cancer

CT imaging data for 117 NSCLC patients were downloaded from the TCIA database and the matched RNA-Seq gene expression profiles were downloaded from the GEO database (GSE103584) [22]. IBSI compliant 397 radiomic features were extracted from ROIs using the CERR radiomics toolbox [23]. A radiomic network was constructed using the extended Bayesian information criterion (EBIC) to optimize the fit of the regularized network constructed using the graphical LASSO [39], resulting in three connected sub-networks that consist of 68 nodes and 88 edges, 56 nodes and 86 edges, and 17 nodes and 26 edges, respectively.

The radio-POSE was constructed as follows: Wasserstein distances were computed between samples over each radiomic network. Similarities corresponding to each radiomic network were computed using ‘k_nn=12’ as the kernel width (Eq. 3). The fused radiomic similarity was then computed as the average similarity and used to compute the DPT distance for constructing the POSE using ‘n_branches=3’.

#### 4.2.4 Glioblastoma multiforme

NetFlow analysis was performed on preprocessed mRNA (*m* = 12,042), DNA methylation (*m* = 1,305) and miRNA expression (*m* = 534) datasets available in the Supplementary Data of [13]. Preprocessing included removal of outliers, imputation for missing data, and normalization. We performed two additional preprocessing steps: first, analysis was restricted to a single sample per patient, resulting in 213 (out of the original 215) samples. Second, we limited the mRNA gene expression analysis to 7,092 genes in the connected HPRD network.

The GBM-POSE was constructed as follows: Euclidean distances were computed between samples for each omics dataset. Similarities for each omics were then computed using ‘k_nn=12’ as the kernel width (Eq. 3) and averaged to construct the fused similarity. The DPT distance was then used as the input to construct the POSE using ‘n_branches=5’.

#### 4.2.5 Simulated datasets

The simulated datasets were generated using the Poisson distribution to create feature profiles with group-specific effects. To simulate a dataset with *m* features, *r* groups, and *n*_*i*_ samples for each group *i*, where *i* = 1, 2, …, *r*, we first generated baseline mean values (*μ*) for each of the *m* features using the Poisson distribution with shape parameter *λ* = 50:

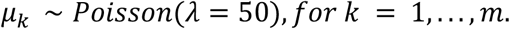

For each sample *j* in some group *i*, the value (*f*_*jk*_) for each feature *k* (where *k* = 1, …, *m*) is sampled from the Poisson distribution with *λ* = *μ*_*k*_ ∗ δ_*jik*_:

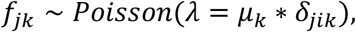

where δ_*jik*_ is the sample’s group-specific offset for feature *k* drawn from the uniform distribution such that

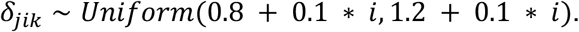

Here, the shape parameter is adjusted by multiplying the baseline feature mean with its corresponding group-specific effect modifier. This approach allowed us to create simulated datasets with non-negative, overlapping but continuous group-specific effects that mimic typical characteristics of high-dimensional biological data.

### 4.3 Benchmark comparison techniques

Comparisons to standard methods were all computed using Python. UMAP was computed with umap-learn [4], t-SNE, LLE, and PCA were computed with scikit-learn [40], and the diffusion technique was computed via scanpy [37].

## Code availability

NetFlow, implemented in Python, is available at https://github.com/areElkin/netflow, with documentation at https://areElkin.github.io/netflow.

## Acknowledgements

This study was supported in part by the NIH/NCI Cancer Center Support grant (grant P30 CA008748), The Simons Foundation, the Breast Cancer Research Foundation (grant MATH-23-001), and the NIH ROBIN cooperative group (grant U54CA274291).

## Contributions

R.E. conceived and developed the approach, implemented the software and performed data analysis. J.H.O. and A.K.S. assisted with data preparation and interpretation of the related analyses. J.O.D. provided project oversight and funding. R.E. wrote the manuscript with input from all authors.

## Competing interests

The authors declare no competing interests.

## Notes

https://github.com/areElkin/netflow

### Competing Interest Statement

The authors have declared no competing interest.

