## Supplementary material for "NetFlow: A Framework to Explore Topological Representations of High-Dimensional Biomedical Data"

Rena Elkin[1], Jung Hun Oh[1], Anish K. Simhal[1], Joseph O. Deasy[1]

[1]*Department of Medical Physics, Memorial Sloan Kettering Cancer Center, New York, NY, 10065, USA*

**Supplementary Algorithm 1:** NetFlow pipeline

**Input:** Dataset $X \in \mathbb{R}^{m \times n}$ with $n$ samples and $m$ features

**Output:** The POSE in the form of a graph $\mathcal{G}_{POSE} = (V, E)$ where vertices correspond to samples

1. Compute distances between samples with metric $d: X \times X \rightarrow \mathbb{R}$
2. Compute affinity matrix $K \in \mathbb{R}^{n \times n}$ from kernel function $k_d: X \times X \rightarrow \mathbb{R}$ where $K_{ij} = k_d(x_i, x_j)$ is computed with respect to the metric $d$.
3. (Optional) Calculate multi-scale or scale-free diffusion distances $\tilde{d}$ using the transition matrix $P \in \mathbb{R}^{n \times n}$ on the weighted graph modeled from the similarity matrix $K$
4. Compute pseudo-ordering backbone using lineage tracing algorithm based on metric, determined by step 1 or optionally computed step 3.
5. Construct the final POSE graph $\mathcal{G}_{POSE}$ by combining nearest neighbor edges to the backbone ordering computed in step 4.

**Supplementary Algorithm 2:** Fused multi-modal/metric NetFlow pipeline

---

**Input:** Datasets $\{X_r\}_{r=1}^{R}$where $X_r \in \mathbb{R}^{m_r \times n}$ with $n$ samples and $m_r$ features

**Output:** The POSE in the form of a graph $\mathcal{G} = (V, E)$ where samples correspond to nodes

1. For each dataset $X_r \in \{K_r\}_{r=1}^{R}$, perform steps 1 and 2 in Algorithm 1 yielding the corresponding set of similarities $\{K_r\}_{r=1}^{R}$
2. Compute the fused similarity: $\widetilde{K} \ = \sum_{r=1}^{R} a_r K_r$ where $\sum_{r=1}^{R} a_r = 1$
3. Continue with step 3 in Algorithm 1, replacing the similarity $K$ with the fused similarity $\widetilde{K}$

---

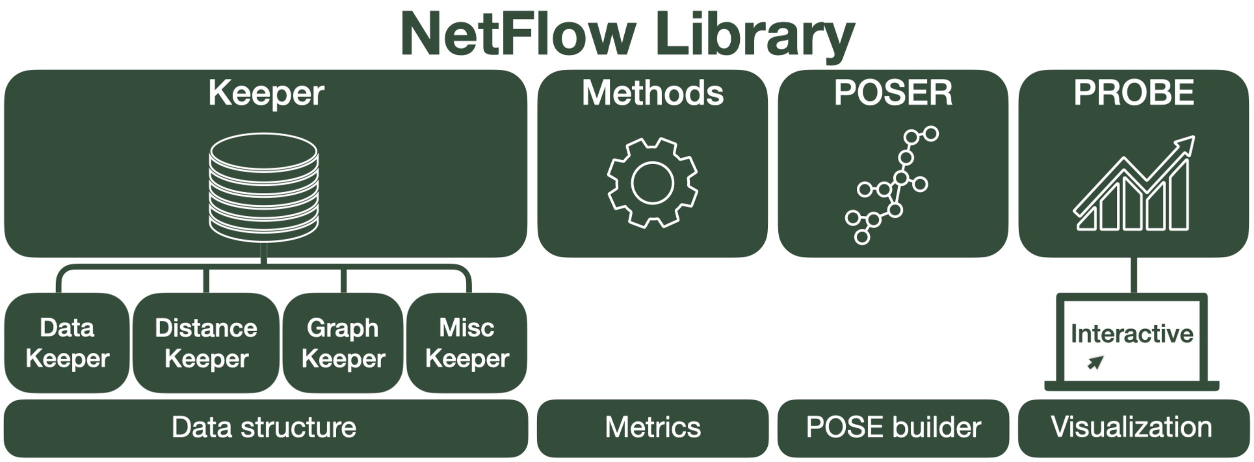


**Supplementary Fig. 1: NetFlow software library.** The NetFlow library consists of the following primary components: Keepers are the data structure used for loading, storing, and saving data. The methods module contains tools for computing metrics and similarities. The POSER module contains tools for computing the POSE representation. The PROBE module contains tools for performing down-stream analysis and visualizing the POSE, including an interactive visualization dashboard. NetFlow is implemented in Python and available at https://areElkin.github.io/netflow.

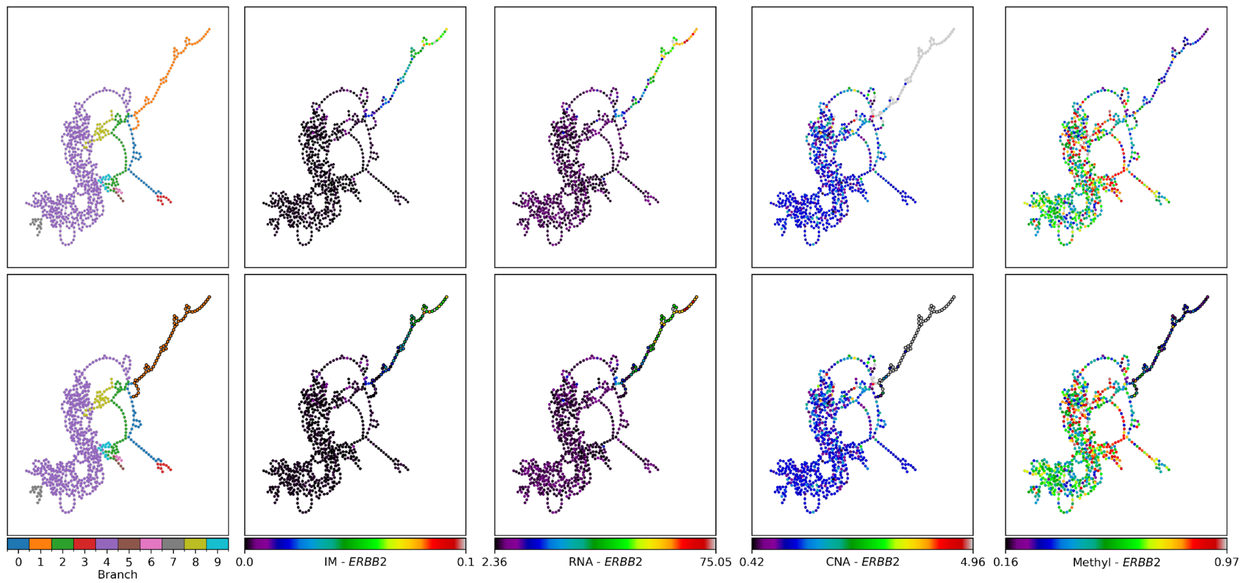


**Supplementary Fig. 2: Branch 1 in the BC-POSE is characterized by ERBB2.** (top) On the left, the BC-POSE is shown with a color overlay annotating the branches. The rest of the subplots, from left to right, are colored by ERBB2 values from the aWCluster multi-omic invariant measure (IM), RNA, CNA, and methylation (methyl) profiles. (bottom) The same is shown below, highlighting branch 1. All samples (nodes) in branch 1 are outlined in black.

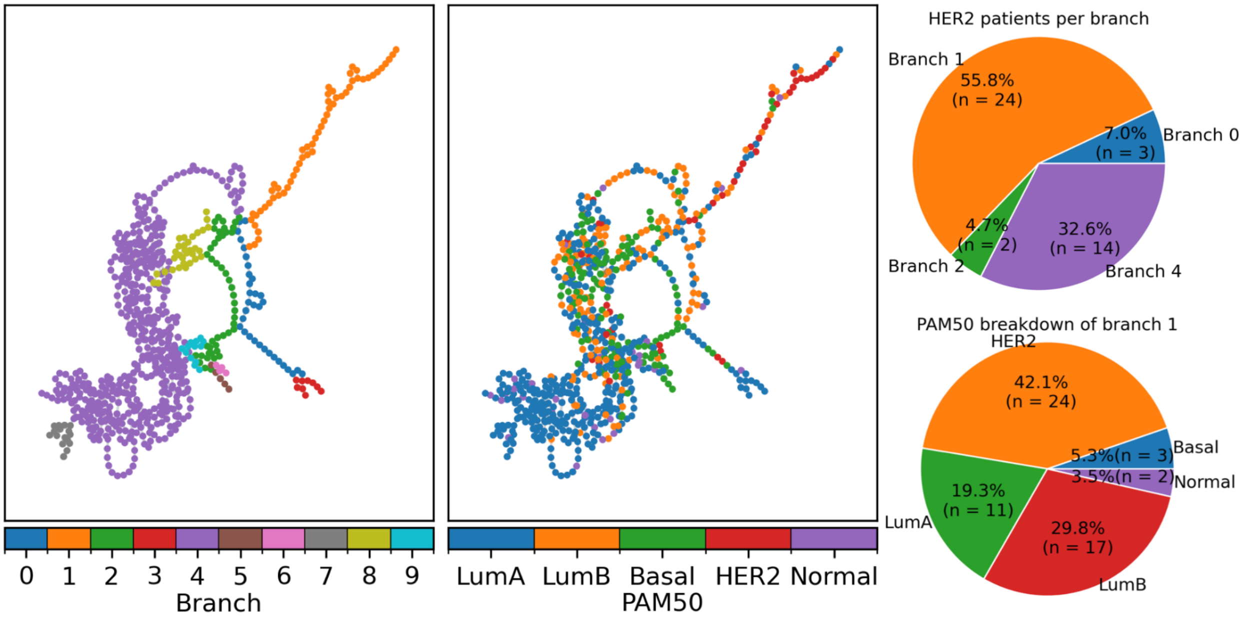


**Supplementary Fig. 3: PAM50 breakdown.** The BC-POSE is color-coded by branch and PAM50 subtypes. Pie charts show (top) percentage of HER2 patients in each branch and (bottom) PAM50 breakdown in branch 1. We find that most HER2 patients are in branch 1 and that branch 1 is predominantly HER2.

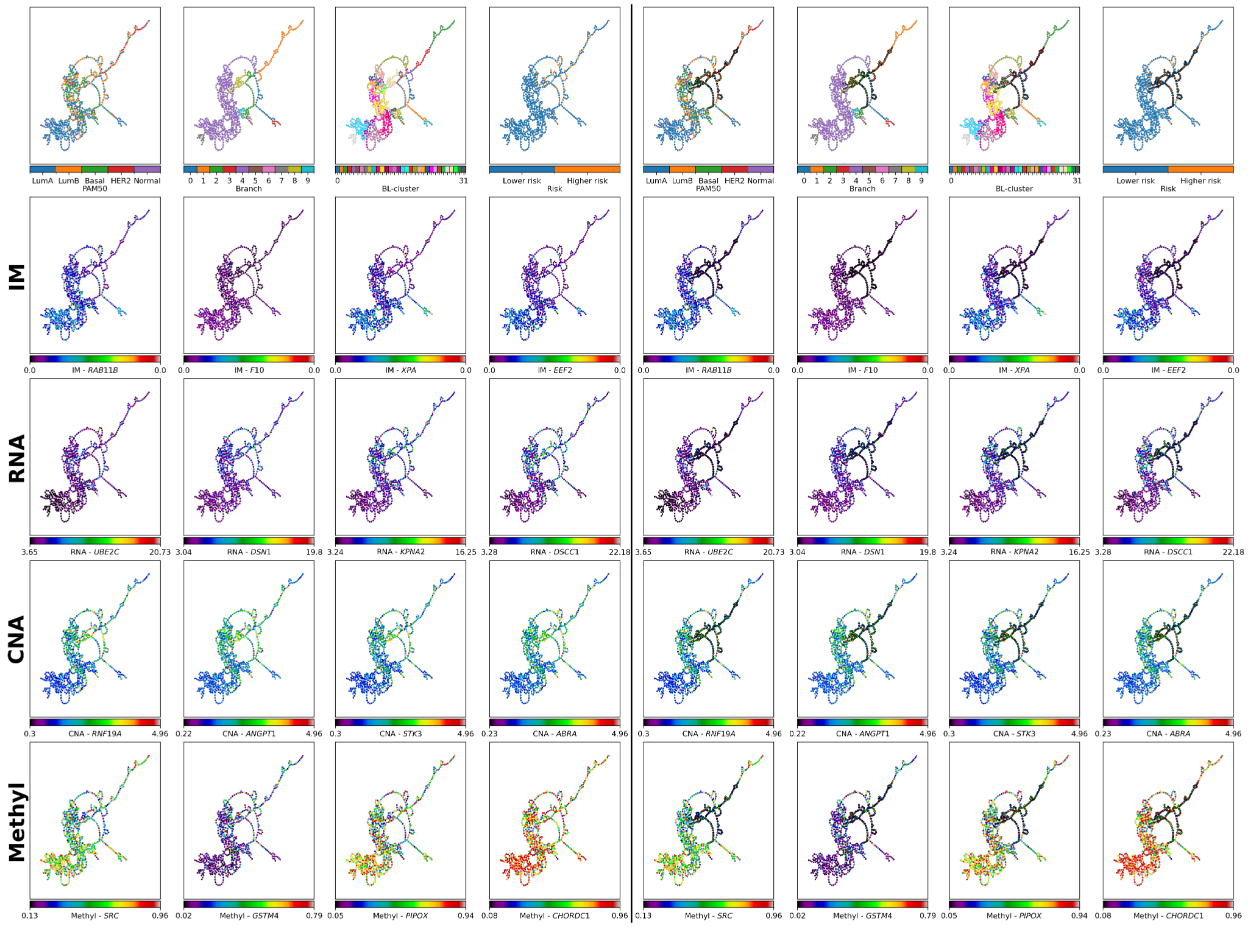


**Supplementary Fig. 4: Characterizing higher-risk BC subtype.** The BC-POSE is shown color-coded by (top row) PAM50 subtype, NetFlow features, and the aWCluster risk group. The rest of the rows, from top to bottom, are colored by the top 4 most significant features associated with the selected samples for the integrated multi-omic invariant measure (IM) profile, RNA, CNA, and methylation (methyl), respectively. The selected samples are outlined in black on the right**.** Tables of top 10 genes significantly associated with the selected samples are shown in Supplementary Tables 1-4.

**Supplementary Table 1: Top significant genes in BC IM data between NetFlow's higher and lower risk cohorts.** Top 10 significant genes with IM profiles differentially associated with the selected samples.

| Feature | High (H) / Low (L) | P value | Corrected P value |
|---|---|---|---|
| *RAB11B* | L | 1.92E-21 | 6.5E-18 |
| *F10* | L | 3.79E-21 | 6.5E-18 |
| *XPA* | L | 7.05E-21 | 8.05E-18 |
| *EEF2* | L | 7.3E-20 | 6.25E-17 |
| *PPM1A* | L | 1.37E-19 | 8.14E-17 |
| *CACNA1C* | L | 1.61E-19 | 8.14E-17 |
| *ZBTB16* | L | 1.66E-19 | 8.14E-17 |
| *PNRC2* | L | 2.65E-19 | 1.13E-16 |
| *NOS2* | H | 3.19E-19 | 1.21E-16 |
| *JDP2* | L | 1.62E-18 | 5.57E-16 |

Mann-Whitney U test for statistical testing with FDR BH correction.

**Supplementary Table 2: Top significant genes in BC RNA data between NetFlow's higher and lower risk cohorts.** Top 10 significant genes with mRNA expression differentially associated with the selected samples.

| Feature | High (H) / Low (L) | P value | Corrected P value |
|---|---|---|---|
| *UBE2C* | H | 4.09E-18 | 8.42E-15 |
| *DSN1* | H | 4.92E-18 | 8.42E-15 |
| *KPNA2* | H | 1.27E-17 | 1.45E-14 |
| *DSCC1* | H | 2.99E-17 | 2.56E-14 |
| *SETBP1* | L | 8.97E-17 | 4.81E-14 |
| *RPN2* | H | 9.27E-17 | 4.81E-14 |
| *BIRC5* | H | 9.82E-17 | 4.81E-14 |
| *EIF2S2* | H | 2.52E-16 | 1.08E-13 |
| *SPAG5* | H | 1.73E-15 | 6.57E-13 |
| *CSE1L* | H | 3.36E-15 | 1.15E-12 |

Mann-Whitney U test for statistical testing with FDR BH correction.

**Supplementary Table 3: Top significant genes in BC CNA data between NetFlow's higher and lower risk cohorts.** Top 10 significant genes with CNA profiles differentially associated with the selected samples.

| Feature | High (H) / Low (L) | P value | Corrected P value |
|---|---|---|---|
| *RNF19A* | H | 9.18E-18 | 2.9E-14 |
| *ANGPT1* | H | 2E-17 | 2.9E-14 |
| *STK3* | H | 2.54E-17 | 2.9E-14 |
| *ABRA* | H | 5.43E-17 | 3.92E-14 |
| *MTSS1* | H | 7.05E-17 | 3.92E-14 |
| *COL14A1* | H | 8.97E-17 | 3.92E-14 |
| *UBR5* | H | 9.12E-17 | 3.92E-14 |
| *RRM2B* | H | 9.15E-17 | 3.92E-14 |
| *LRP12* | H | 1.23E-16 | 4.67E-14 |
| *DSCC1* | H | 1.53E-16 | 4.72E-14 |

Mann-Whitney U test for statistical testing with FDR BH correction.

**Supplementary Table 4: Top significant genes in BC methylation data between NetFlow's higher and lower risk cohorts.** Top 10 significant genes with methylation profiles differentially associated with the selected samples.

| Feature | High (H) / Low (L) | P value | Corrected P value |
|---|---|---|---|
| *SRC* | L | 1.56E-14 | 5.33E-11 |
| *GSTM4* | H | 3.08E-11 | 5.28E-08 |
| *PIPOX* | L | 1.66E-10 | 1.89E-07 |
| *CHORDC1* | L | 6.65E-10 | 5.69E-07 |
| *TNFSF12* | H | 1.48E-09 | 1.01E-06 |
| *HSP90AA1* | L | 1.86E-09 | 1.06E-06 |
| *BNIP3* | L | 3.26E-09 | 1.59E-06 |
| *AANAT* | H | 4.19E-09 | 1.8E-06 |
| *RRM2B* | L | 6.07E-09 | 2.28E-06 |
| *LGALS3* | H | 7.07E-09 | 2.28E-06 |

Mann-Whitney U test for statistical testing with FDR BH correction.

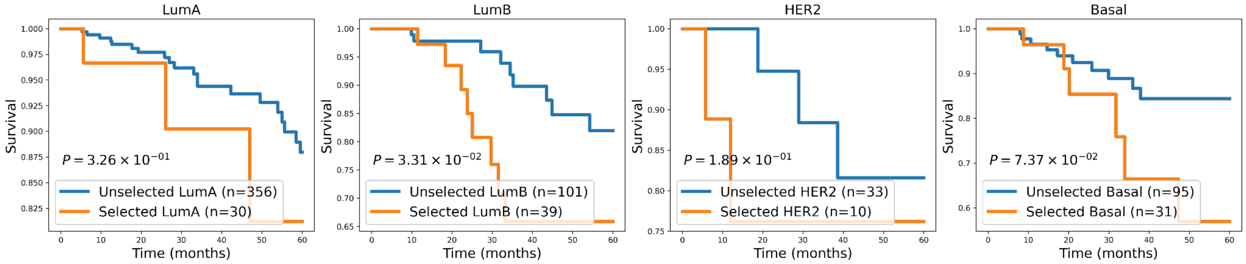


**Supplementary Fig. 5: PAM50 subtype inspection of BC-POSE selected higher-risk samples.** Kaplan-Meier survival curve analysis for each PAM50 subtype between selected and not selected samples from the BC-POSE (logrank test, $P$ values are shown).

**Supplementary Table 5: Differences in CoMMpass data patient characteristics between NetFlow's *WEE1*-based higher and lower risk cohorts.**

| Feature name | Low-risk cohort (n=490) | High-risk cohort (n=169) | P value |
|---|---|---|---|
| Age (mean ± S.D.) | 62.9 ± 10.2 | 61.6 ± 11.9 | 0.176 |
| Sex | M: 282; F: 208 | M: 111; F: 58 | 0.0691 |
| ISS | (1): 178; (2): 174; (3): 124 | (1): 51; (2): 58; (3): 55 | 0.146 |
| R-ISS | I: 29.2%; II: 62.8%; III: 8% | I: 17.7%; II: 67.7%; III: 14.6% | 2.26E-3 |
| Hyperdiploidy | 63.1% | 30.2% | 1.30E-13 |
| t(4;14) | 13.3% | 11.8% | 0.730 |
| t(11;14) | 13.1% | 40.2% | 6.38E-14 |
| MAF translocation | 2.86% | 7.10% | 0.0268 |
| MYC translocation | 15.3% | 7.10% | 9.72E-3 |
| Hyper APOBEC | 4.38% | 14.8% | 8.44E-5 |
| Gain 1q21 | 33.1% | 39.6% | 0.0762 |
| TP53 mutation | 6.30% | 18.9% | 1.35E-6 |
| Treatment | combined bortezomib/IMIDs-based: 234; Bortezomib-based: 100; combined IMIDs/carfilzomib-based: 87; IMIDs-based: 25; Carfilzomib-based: 24; combined bortezomib/IMIDs/carfilzomib-based: 19; combined bortezomib/carfilzomib-based: 1; | combined bortezomib/IMIDs-based: 83; Bortezomib-based: 38; combined IMIDs/carfilzomib-based: 23; Carfilzomib-based: 14; IMIDs-based: 7; combined bortezomib/IMIDs/carfilzomib-based: 4; | - |

S.D.: standard deviation, ISS: International Staging System, R-ISS: Revised ISS. Chi-Squared test for statistical testing, except for 'age,' tested via t-test.

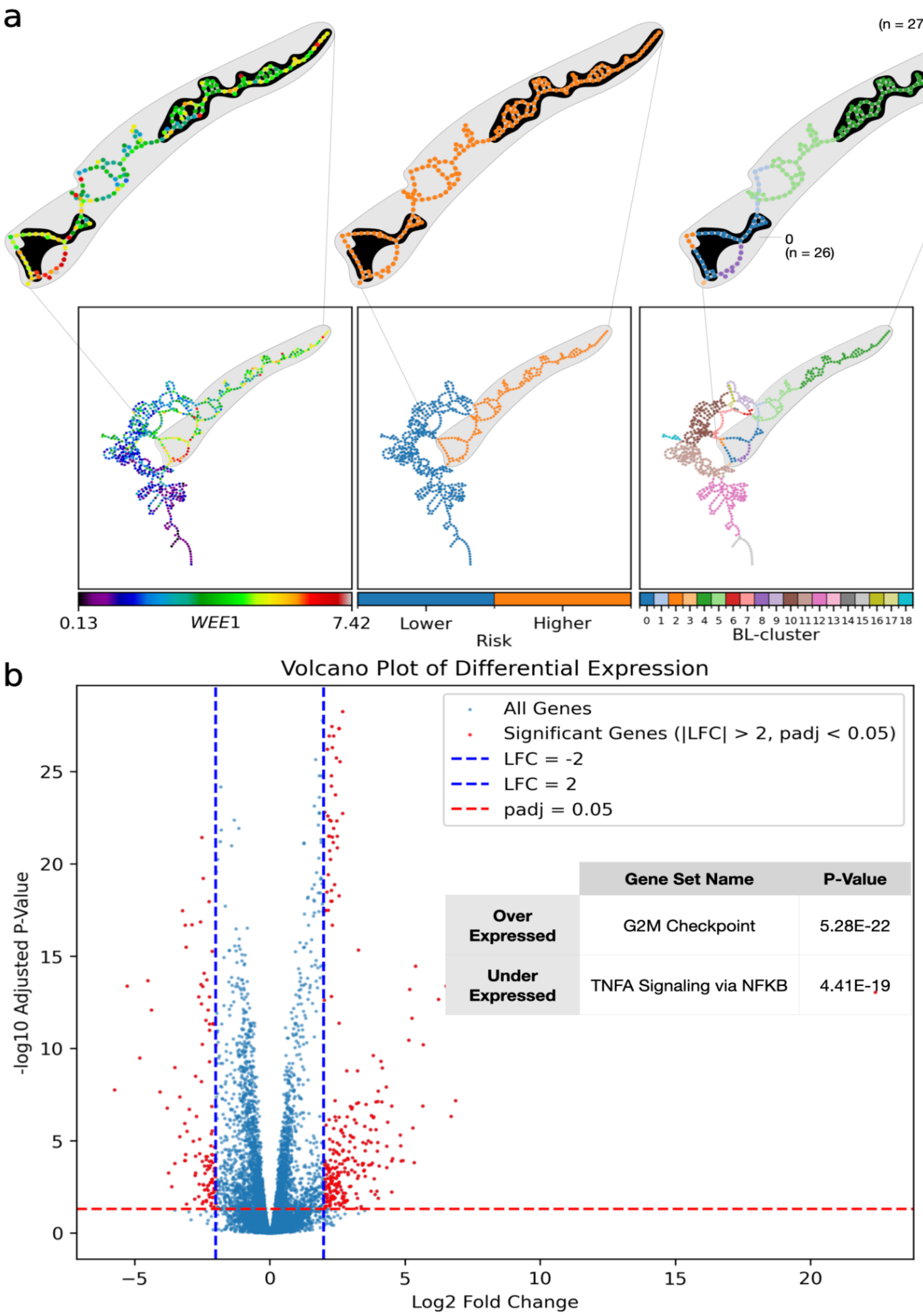


**Supplementary Fig. 6: Varied implications of high *WEE1* expression.** (a) Comparison of BL-clusters on opposite ends (right) of identified higher-risk group MM-POSE (middle), associated with high *WEE1* expression (left). (b) Volcano plot of differential gene expression between accented BL-cluster clusters 0 (baseline) and 4. Top significant pathways associated with over- and under-expressed genes are shown along with FDR corrected $P$ values. Gene set enrichment analysis was performed via mSigDB using the GSEA hallmark pathways database [1-2].

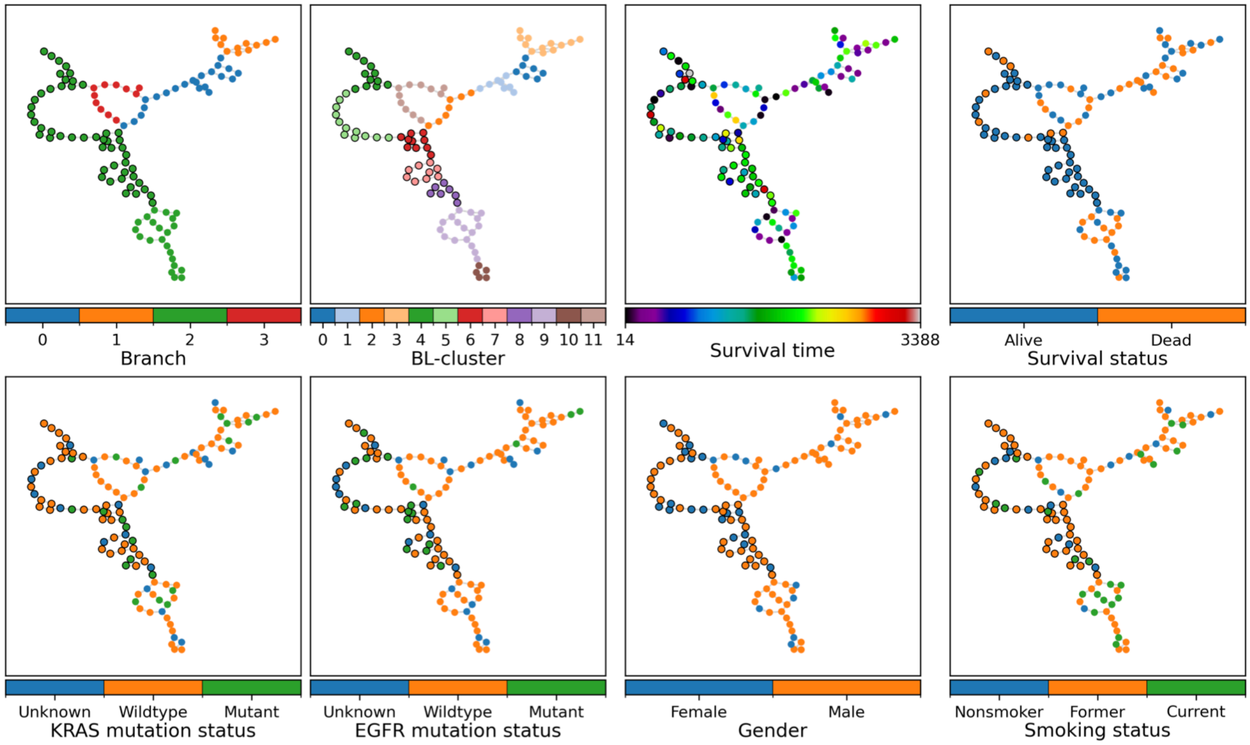


**Supplementary Fig. 7: The radio-POSE**. Each node represents a sample and the nodes corresponding to selected samples are shown outlined in black. Nodes are colored by NetFlow and clinical features.

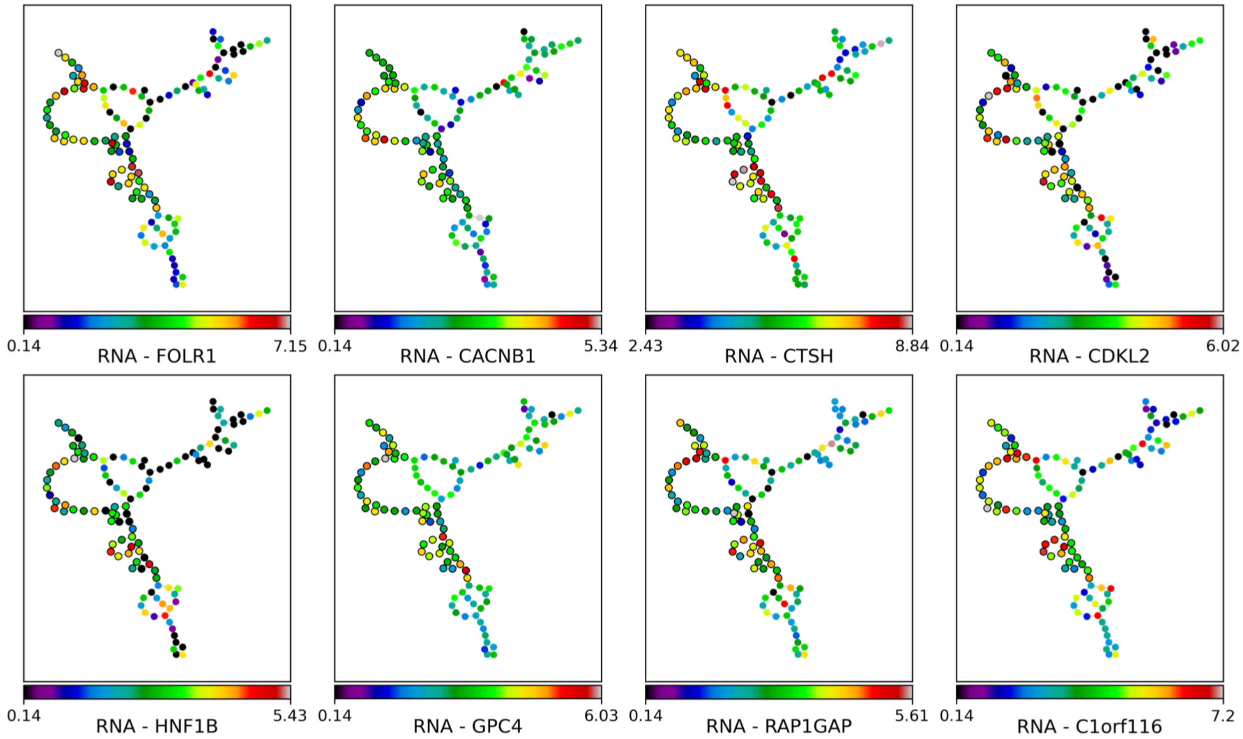


**Supplementary Fig. 8: Genes significantly associated with selected samples in the radio-POSE**. Each node represents a sample and the nodes corresponding to selected samples are shown outlined in black. Nodes are colored by the eight significant genomic features associated with the selected samples (Mann-Whitney U test with FDR-BH correction; $P$ values: *FOLR1*=0.01; *CACNB1*=0.03; *CTSH*=0.04; *CDKL2*=0.04; *HNF1B*=0.04; *GPC4*=0.04; *RAP1GAP*=0.04; *C1orf116*=0.04).

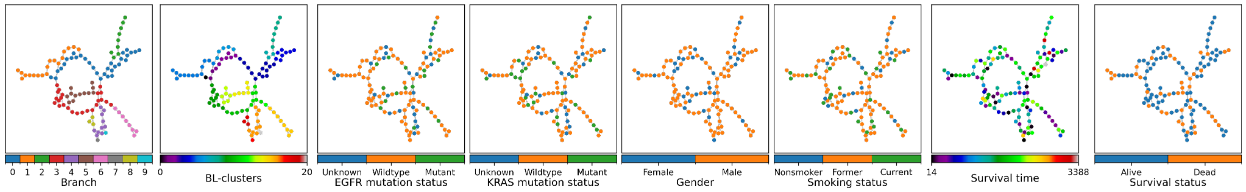


**Supplementary Fig. 9: radio-POSE based on Euclidean distances of radiomic features do not associate with genomic features.** NSCLC fused radio-POSE based off of the Euclidean distance

**Supplementary Table 6: Top significant characteristics associated with GBM lower-risk BL-cluster 17 subtype.**

| | Feature | High (H) / Low (L) | P value | Corrected P value |
|---|---|---|---|---|
| RNA | *EMP3* | L | 1.63e-12 | 5.77e-9 |
| | *TIMP1* | L | 2.41e-12 | 5.77e-9 |
| | *EFEMP2* | L | 2.97e-12 | 5.77e-9 |
| | *PLA2G2A* | L | 4.86e-12 | 5.77e-9 |
| | *MAPK8* | H | 5.24e-12 | 5.77e-9 |
| Methyl | *CRIP1* | H | 2.51e-9 | 3.27e-6 |
| | *FES* | H | 4.65e-8 | 1.89e-5 |
| | *IL17RB* | H | 5.01e-8 | 1.89e-5 |
| | *RAB32* | H | 5.8e-8 | 1.89e-5 |
| | *PAX6* | H | 8.25e-8 | 2.15e-5 |
| miRNA | *miR-222* | L | 1.99e-10 | 6.59e-8 |
| | *miR-221* | L | 2.47e-10 | 6.59e-8 |
| | *miR-34a* | L | 3.08e-9 | 5.49e-7 |
| | *miR-34b* | L | 1.75e-8 | 2.34e-6 |
| | *miR-328* | H | 2.4e-8 | 2.57e-6 |

Mann-Whitney U test for statistical testing with FDR BH correction.

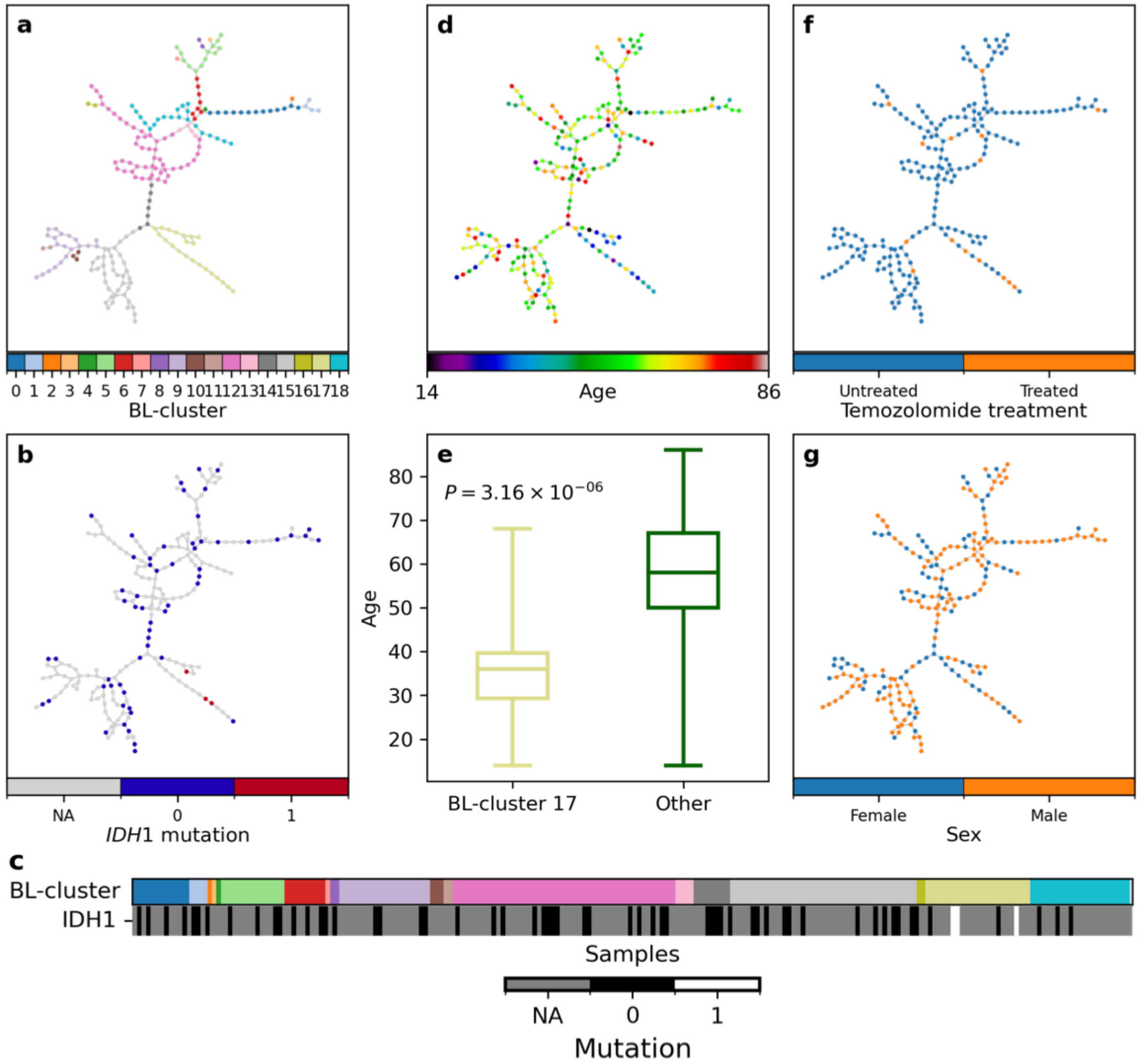


**Supplementary Fig. 10: Clinical characteristics of higher-risk BL-cluster 17 subgroup.** (a) The GBM-POSE is displayed with nodes colored by their BL-cluster membership (the same as shown in Fig. 5a). (b) IDH1 mutation status is shown on the GBM-POSE and (c) by BL-cluster. Of the samples with mutational data, only a few samples were found to have an IDH1 mutation, however all those that did were found in BL-cluster 17. (d) looking at age (years), we find that patients whose samples are in the lower-risk BL-cluster 17 subtype tend to be younger. (e) Box plots of age distribution for patients in the lower-risk BL-cluster 17 and all other patients (boxes span the first and third quartiles with a line at the median, and the whiskers cover the whole range of age values; Welch's two-sided t-test $P$ value is shown). (f) POSE colored by treatment status (treated vs untreated in BL-cluster 17: logrank p = 0.88). (g) POSE colored by sex (BL-cluster 17 vs other: Fisher's exact test, $P = 0.24$). These findings are consistent with the lower-risk group identified in the original Similarity Network Fusion (SNF) paper [13], which was characterized by IDH1 mutation and tended to be younger in age. Also consistent with the SNF lower-risk group, no significant difference in terms of survival was found between treated vs untreated samples.

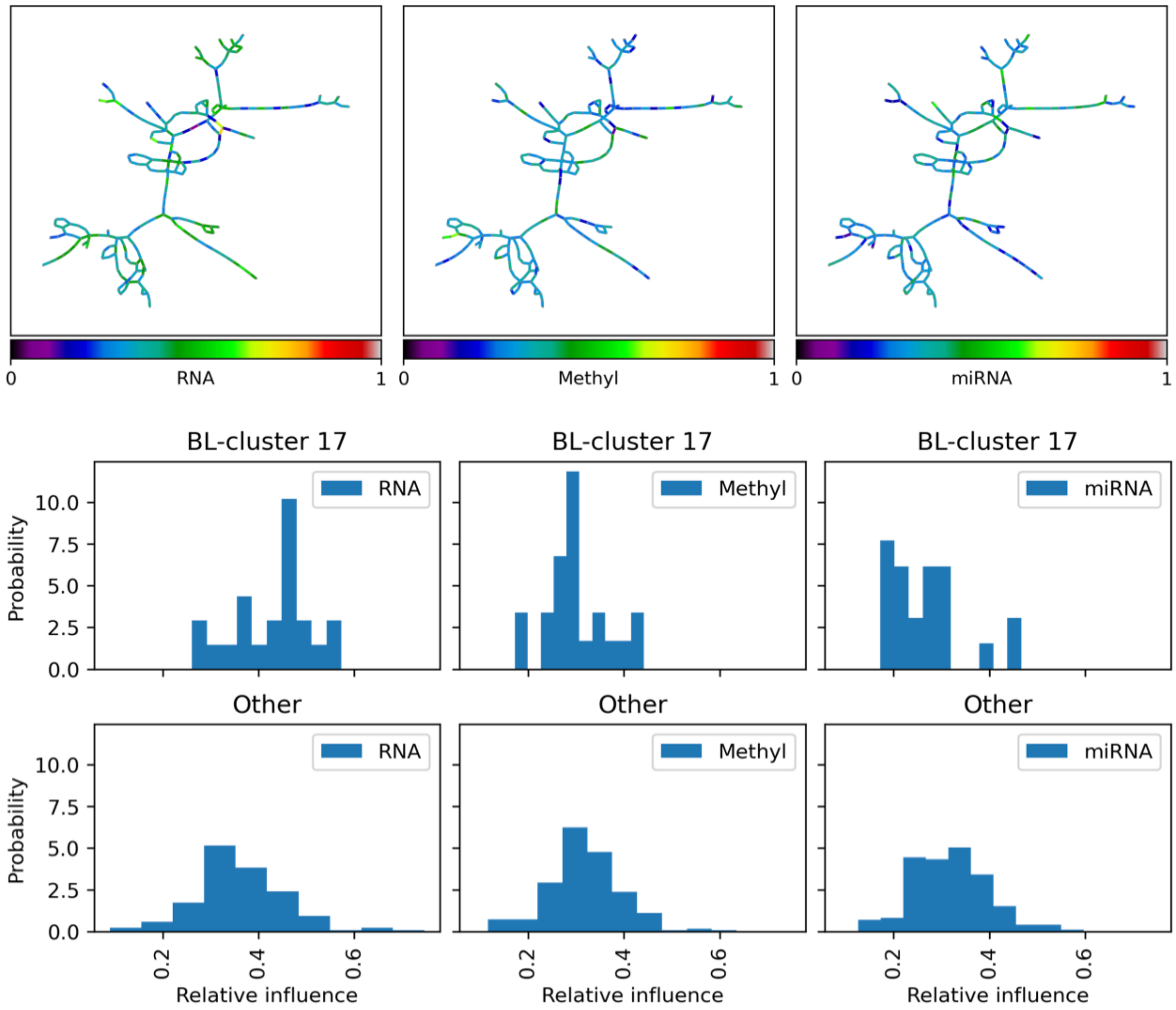


**Supplementary Fig. 11: Relative influence of omics modality on GBM-POSE edges.** (top) Edges in the GMB-POSE are colored by the strength of each data modality's relative influence. The relative influence of a modality on an edge is considered to be the transition probability between associated samples according to the diffusion operator P for that modality normalized by that edge's net transition probability summed over all modalities. (bottom) Histograms of relative edge influence of each modality for edges within BL-cluster 17 and the other samples. Overall, the distributions of each modality's influence are consistent between edges within BL-cluster 17 and the rest of the edges.

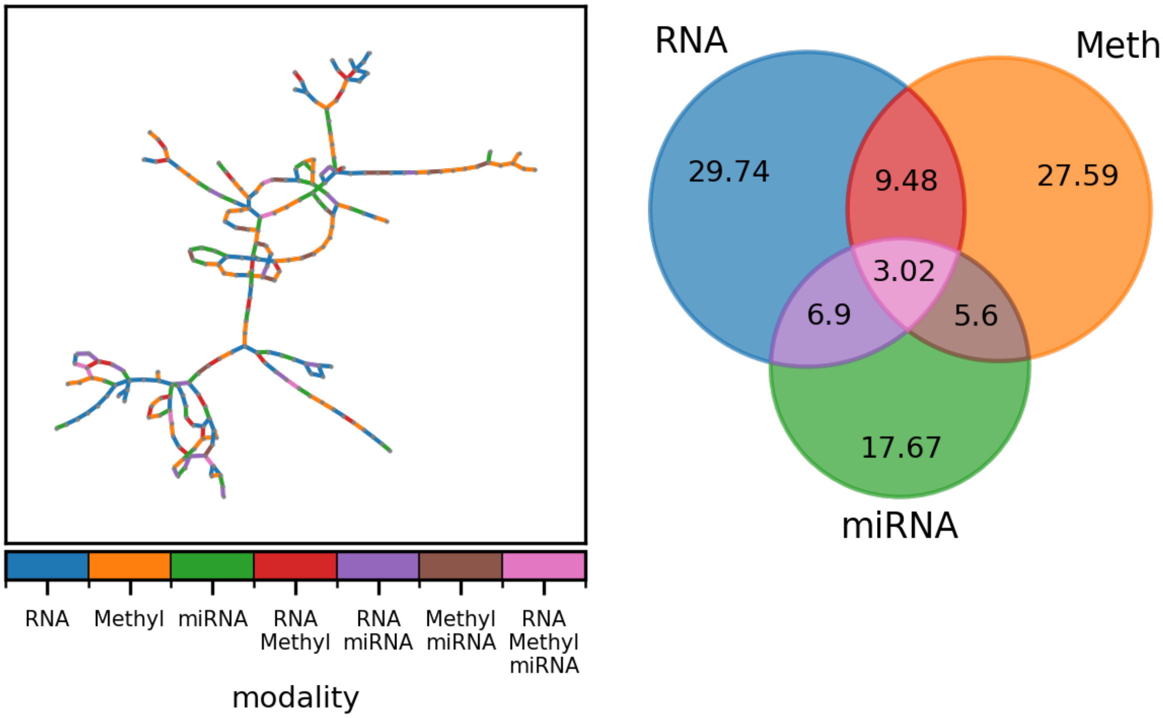


**Supplementary Fig. 12. Data source attributions of GBM-POSE edges.** (left) GBM-POSE with edges colored by attributed source data modality. (right) Venn diagram of edge source associations by data modality (%). Edges were attributed to data modalities, as described in the original SNF paper [13]. Briefly, an edge is attributed to a single modality if the similarity with respect to that modality was at least 10% higher than the similarity in any other modality. If the difference in similarity between the two highest modalities is less than 10%, the edge is attributed to both modalities, and if the difference in similarities between all modalities is less than 10%, the edge is attributed to all modalities. The Venn diagram of the percentage of source modalities attributed to each edge shows that no single modality independently characterizes the POSE.

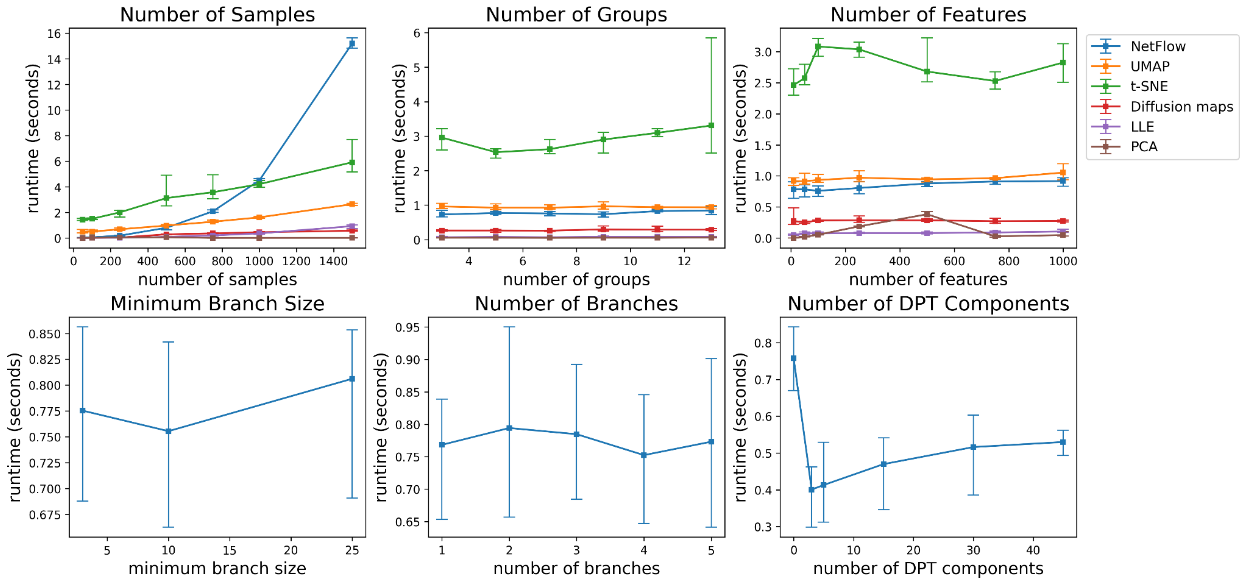


**Supplementary Fig. 13: Assessing parameter effects on NetFlow runtime in comparison to benchmark approaches.** (Top) When simulating the data, we used a fixed default value for each variable. Considering that UMAP usage is recommended for datasets with at least 500 samples, we set the default number of samples to 500. For the other parameters, we set the default number of groups to be 3 and the default number of features to be 100. To provide information on how the runtime scales with each variable independently, we then perturbed one variable at a time while the other variables were kept fixed according to their default values. To this end, we simulated datasets iterating the number of samples over (50, 100, 250, 500, 750, 1000, 1500), the number of groups was iterated over (3, 5, 7, 9, 11, 13), and the number of features was iterated over (10, 50, 100, 250, 500, 750, 1000). Each combination of variable values is simulated 5 times, resulting in a total of 100 generated datasets. (Bottom) We also consider how the choices of NetFlow-specific parameters affect runtime. Specifically, we consider the following parameters: ‘min_branch_size’ with a default value of 10, ‘n_branches’ with a default value of 3, and ‘n_dpt_comps’ with a default value of 0. Following the DPT branching algorithm, as implemented in the Python ScanPy library, the minimum branch size is used to specify the minimum number of samples that should remain in a branch and are therefore not considered as a possible branch point. The number of branches specifies the number of times branching should be iteratively attempted and the number of DPT components specifies the number of diffusion components to be used when computing the multi-scale DPT distance. When the number of DPT components is set to 0, all diffusion components are used. As above, the parameters (including the variables assessed above) are set to the default values, and each NetFlow-specific parameter is perturbed at a time. We iterate minimum branch size over (3, 10, 25), number of branches over (1, 2, 3, 4, 5), and the number of DPT components over (3, 5, 15, 30, 50, 100, 250, 500). As before, each combination of parameters was simulated 5 times, resulting in 80 simulations. We see that the minimum branch size and number of branches have little effect on runtime. Reducing the number of DPT components, especially when the number of samples is large, may be preferential to improve computational efficiency, but at the cost of precision.
